# A synaptic novelty signal to switch hippocampal attractor networks from generalization to discrimination

**DOI:** 10.1101/2021.02.24.432612

**Authors:** Ruy Gómez-Ocádiz, Massimiliano Trippa, Lorenzo Posani, Simona Cocco, Rémi Monasson, Christoph Schmidt-Hieber

## Abstract

Episodic memory formation and recall are complementary processes that put conflicting requirements on neuronal computations in the hippocampus. How this challenge is resolved in hippocampal circuits is unclear. To address this question, we obtained *in vivo* whole-cell patch-clamp recordings from dentate gyrus granule cells in head-fixed mice trained to explore and distinguish between familiar and novel virtual environments. We find that granule cells consistently show a small transient depolarization of their membrane potential upon transition to a novel environment. This synaptic novelty signal is sensitive to local application of atropine, indicating that it depends on metabotropic acetylcholine receptors. A computational model suggests that the observed transient synaptic response to novel environments may lead to a bias in the granule cell population activity, which can in turn drive the downstream attractor networks to a new state, thereby favoring the switch from generalization to discrimination when faced with novelty. Such a novelty-driven cholinergic switch may enable flexible encoding of new memories while preserving stable retrieval of familiar ones.

## Introduction

The hippocampus is essential for the encoding, storage and retrieval of episodic memories^1–4^. Recall of these memories is thought to result from the reactivation of previously stored patterns of neural activity in the hippocampal regions CA3 and CA1, even when the inputs from upstream circuits are degraded or incomplete. Such neuronal generalization (‘pattern completion’) is supported by attractor network dynamics arising from recurrent connectivity among pyramidal cells in these hippocampal regions, which enables the network to reinstate the activity of previously established neuronal assemblies^2,4–9^. A complementary neuronal discrimination (‘pattern separation’) process needs to be engaged if the differences between ongoing experiences and previously stored representations exceed a threshold for behavioral relevance, requiring the hippocampus to encode and store novel episodic memories in new cell assemblies. However, memory formation and recall put conflicting requirements on hippocampal computations, as the reliable retrieval of familiar representations supported by robust attractor properties in the CA3 and CA1 circuits opposes the formation of new neuronal assemblies for the storage of novel episodic memories. How the hippocampal network reconciles these conflicting demands to achieve an optimal balance between memory formation and recall remains unclear^10–12^.

The dentate gyrus, situated immediately upstream of the CA3 region, appears well suited to solve this problem, as it performs neuronal discrimination by orthogonalizing multimodal inputs from the entorhinal cortex through sparse firing activity and cellular expansion^2,6–8,12–21^. Hence, the dentate gyrus could be charged with the task of detecting novelty and selectively reporting it to downstream circuits, instructing them to store a new representation through a shift towards a different attractor state. However, experimental data have shown that the dentate gyrus robustly reports differences between any environments, independent of whether they are novel or familiar^12^.

Several requirements on the hippocampal memory system can explain that the dentate gyrus acts as a neutral difference detector. First, some aspects of a given experience might be categorized as familiar and thus lead to recall, while others are identified as novel and thereby favor encoding of a new memory. For example, when encountering a familiar location, it is equally important to retrieve the corresponding memory through generalization as it is to detect the differences between the present episode and the memorized representation through discrimination to update memory with the ongoing experience^21–23^. Therefore, decorrelated outputs from the dentate gyrus need to be able to simultaneously support recall of familiar representations and drive the formation of new representations in downstream regions. Moreover, whether a novel experience is sufficiently behaviorally relevant to merit encoding as a separate memory depends on the current behavioral context, general alertness and arousal state, which are typically conveyed by extrahippocampal signals^22,24–26^. Given that the dentate gyrus performs neuronal discrimination steadily, regardless of whether the ongoing experience is novel or familiar, how can its robust discrimination code be selectively used to direct downstream circuits to new attractor states that encode new episodic memories?

To address this question, we obtained *in vivo* whole-cell patch-clamp recordings from dentate gyrus granule cells in head-fixed mice trained to explore and distinguish between familiar and novel virtual environments^12,27^. We report that granule cells consistently show a task-dependent small and transient depolarization of their membrane potential when an animal encounters a novel environment. This depolarization can be abolished by local application of atropine, indicating that it depends on metabotropic acetylcholine receptors. A computational model suggests that the observed synaptic response to environmental novelty leads to a bias in the granule cell population activity which can drive the CA3 attractor network to a new state, thereby favoring discrimination during memory encoding, as opposed to the default generalization underlying recall when the animal navigates in a familiar environment. Our experimental results and our model can explain how an external cholinergic signal enables the hippocampus to effectively encode novel memories while preserving stable retrieval of familiar ones.

## Results

### Mice can distinguish between familiar and novel virtual reality environments

To explore the behavioral effect of novelty, we used an immersive virtual reality setup adapted for rodent head-fixed navigation (Fig. 1a). We created three visually-rich virtual reality environments with identical geometry and task logic but with different sets of proximal and distal cues^28^ and wall and floor textures. After habituation to the virtual reality setup, we trained water-restricted mice to navigate in the virtual corridor of the familiar environment (F) and stop at an un-cued reward zone for a defined period of time to obtain a water reward. Mice were ‘teleported’ back to the beginning upon arrival at the end of the corridor (Fig. 1b). We observed a marked increase in performance across five consecutive daily training sessions in the familiar environment (F) of 20-30 min each, as measured by licking in anticipation of reward delivery (hit rate: 0.03 ± 0.01 hits/lap during the 1st session and 0.25 ± 0.04 hits/lap during the 5th session; *n* = 57 mice, paired t-test, *p* < 0.001), reflecting the reliable learning of the task (Fig. 1c,d). On the sixth training session, we introduced the novel environment 1 (N1) and compared the performance in both environments. We found that while the performance in the familiar environment (F) reflected the learning of the task, the performance in the novel environment 1 (N1) decreased to a level comparable to untrained animals (hit rate: 0.21 ± 0.05 hits/lap in F and 0.06 ± 0.03 hits/lap in N1; *n* = 57 mice, paired t-test, *p* < 0.001; first session in F versus N1, paired t-test, *p* > 0.05; Fig. 1d). These results indicate that mice can rapidly learn to perform the virtual reality navigation task and that this learning is environment-specific, as evidenced by the ability of the animals to show different behavior when exposed to novelty.

**Fig. 1.**
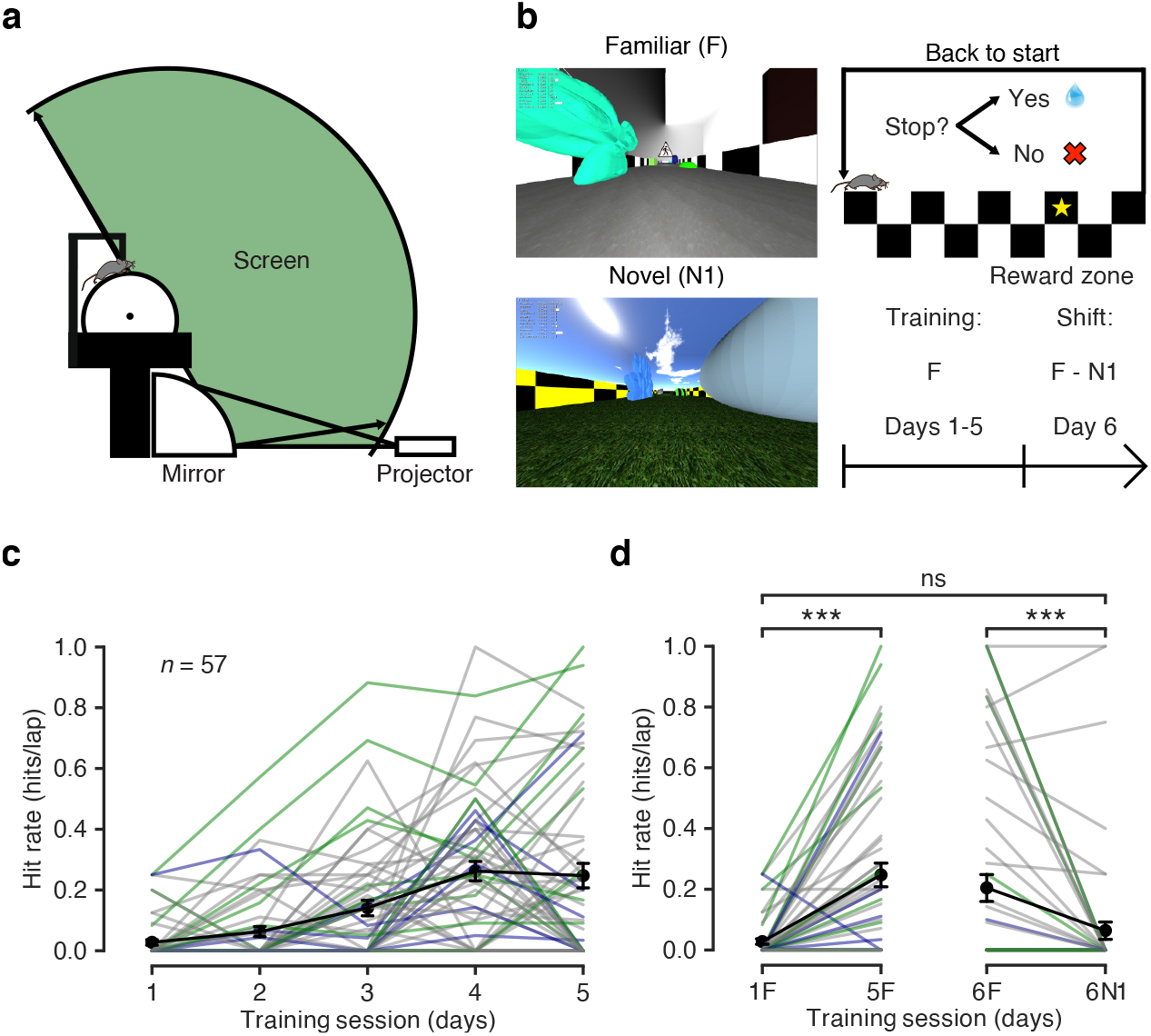
Mice can distinguish between familiar and novel virtual reality environments. **a** Schematic of the virtual reality setup (adapted from Schmidt-Hieber & Häusser, 2013). **b** Left: view along the long axis of the corridor in the familiar (F, top) and the novel 1 (N1, bottom) virtual reality environments. Top right: schematic of the navigation task. Animals are trained to stop in a reward zone to obtain a reward. At the end of the corridor, animals are ‘teleported’ back to the beginning. Bottom right: behavioral protocol timeline. **c** Behavioral performance across training sessions, measured as hit rate (see Methods). Grey lines represent individual animals. Animals included in the experiments from Fig. 2 and Fig. 4 are highlighted in green (controls) and mauve (local application of atropine). Black circles represent the mean ± s.e.m. across all animals (*n* = 57 mice). **d** Left: comparison of behavioral performance during the first (1F) and the last (5F) training sessions in the familiar environment (hit rate: 0.03 ± 0.01 hits/lap and 0.25 ± 0.04 hits/lap, respectively; *n* = 57 mice, paired t-test, *p* < 0.001). Right: comparison of behavioral performance in the familiar (6F) and in the novel environment 1 (6N1) during the 6th training session (hit rate: 0.21 ± 0.05 hits/lap and 0.06 ± 0.03 hits/lap, respectively; *n* = 57 mice, paired t-test, *p* < 0.001; 1F versus 6N1, paired t-test, *p* > 0.05).

### Granule cells transiently depolarize in response to saliency

To study the synaptic integration processes underlying this behavioral discrimination^29,30^, we obtained whole-cell patch-clamp recordings from granule cells while mice performed the task, but this time switching between laps in the familiar environment (F) and in the novel environment 2 (N2) to ensure that the animals had never encountered the novel environment before (Fig. 2a,b). We restricted our experiments to one recorded cell per animal and we confirmed the accuracy of the recording site in all cases using electrophysiological and anatomical criteria (see Methods). We then computed the teleportation-aligned average of the membrane potential traces around the teleportation events and compared the teleportations within the familiar environment (FF) and between the familiar environment and the novel environment 2 (FN2) (Fig. 2c-e). We did not observe any significant difference between the overall mean membrane potential recorded during the total time spent in the familiar and novel environment (−68.9 ± 4.1 mV in F and −68.9 ± 4.1 mV in N2; *n* = 9 cells, paired t-test, *p* > 0.05; Fig. 2e). However, we found a consistent membrane potential depolarization following the teleportation event when the novel environment 2 (N2) was introduced for the first time (mean Δ*V*_m_ during 1 s after the teleportation: 1.02 ± 0.25 mV for FN2 teleportations versus −0.01 ± 0.16 mV for FF teleportations; *n* = 9 cells, paired t-test, *p* < 0.01; Fig. 2e and Fig. S1b). This *V*_m_ depolarization was not explained by a change in animal movement upon encountering the novel environment, as we did not observe a significant difference in speed between FF and FN2 teleportations (Δspeed: 0.0 ± 0.9 cm/s for FN2 versus −0.8 ± 0.4 cm/s for FF; *n* = 9 cells, paired t-test, *p* > 0.05; Fig. S1c). The depolarization lasted for ~2 s (see Fig. S1a), was not accompanied by a change in membrane potential variance (1.4 ± 1.0 mV^2^ for FN2 teleportations versus −0.5 ± 0.4 mV^2^ for FF teleportations; *n* = 9 cells, paired t-test, *p* > 0.05; Fig. 2e) and its magnitude was not correlated with the mean membrane potential of the cells (FN2 Δ*V*_m_ versus mean *V*_m_: Pearson’s correlation coefficient, *r* = −0.19, *p* > 0.05; Fig. S1d). Since these recordings were obtained using a blind unbiased approach, the consistent observation of a transient membrane potential depolarization suggests that this synaptic phenomenon is not restricted to a specific subset of dentate gyrus neurons.

**Fig. 2.**
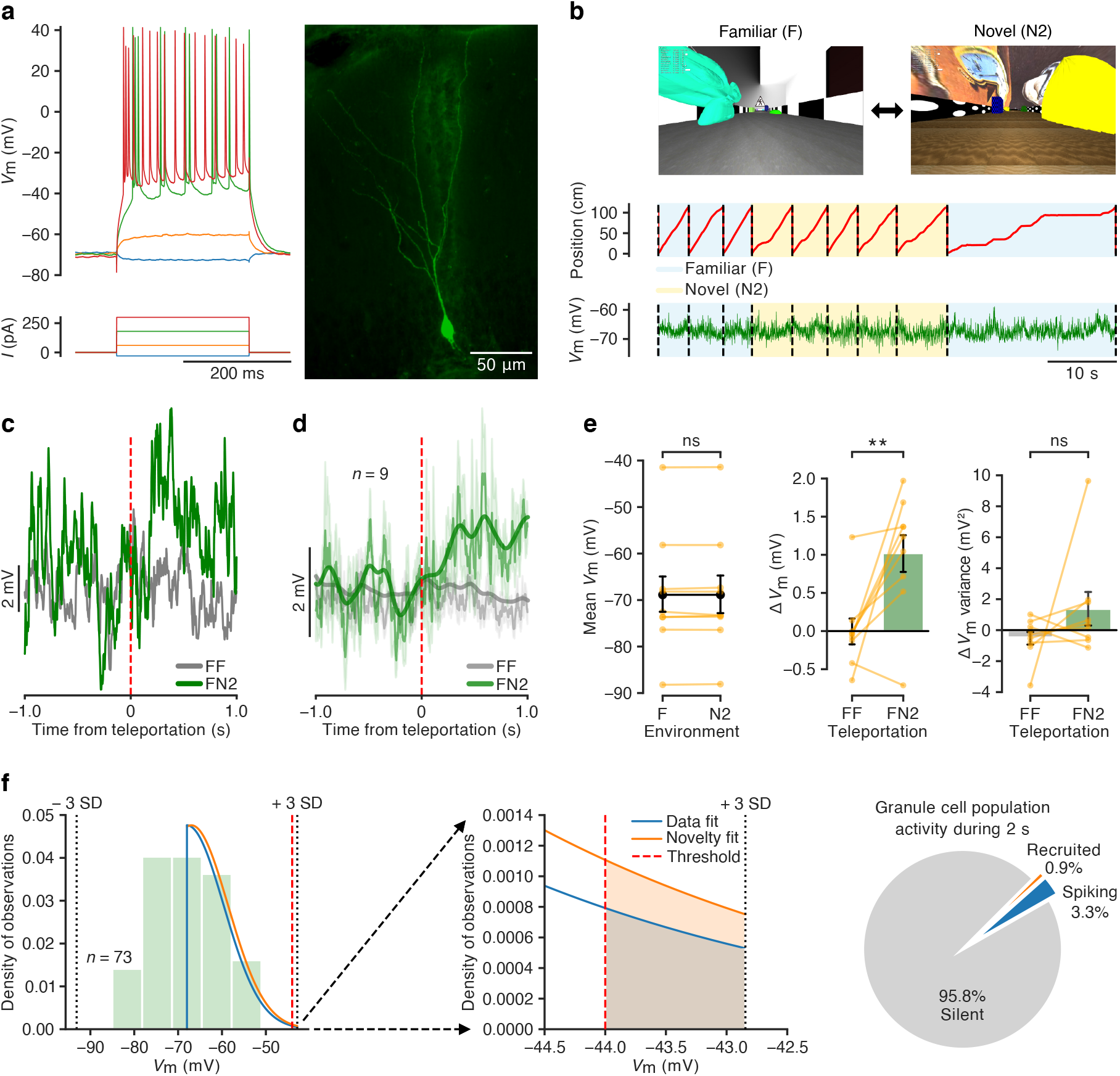
Granule cells transiently depolarize in response to saliency. **a** Example recording from a granule cell. Left: membrane potential responses to current pulse injections. Right: fluorescence image obtained after biocytin filling during the recording. **b** Example recording during a behavioral discrimination experiment switching between the familiar (F) and a novel environment (N2). Top: view along the long axis of the corridor in the familiar (F, left) and the novel 2 (N2, right) environments. Traces show animal position along the corridor (middle) and *V*_m_ (bottom). **c** Teleportation-aligned average from a representative recording. Traces represent the mean *V*_m_ aligned to the teleportation events. The average from teleportations within the familiar environment (FF) is shown in grey and the teleportation from the familiar to the novel environment (FN2) is shown in green. Teleportation time is indicated by the vertical red dashed line. **d** Teleportation-aligned average across multiple recordings. Traces represent the mean ± s.e.m. of the *V*_m_ aligned to the teleportation events. Teleportations within the familiar environment (FF) are shown in grey and teleportations from the familiar to the novel environment (FN2) are shown in green. Teleportation time is indicated by the vertical red dashed line. Low-pass filtered traces (bold traces) are shown superimposed. **e** Left: summary of mean *V*_m_ for the familiar (F) and the novel (N2) environments (−68.9 ± 4.1 mV and 68.9 ± 4.1 mV, respectively; *n* = 9 cells, paired t-test, *p* > 0.05). Middle: Δ*V*_m_ summary for the teleportations within the familiar environment (FF) and the teleportations from the familiar to the novel environment (FN2) (−0.01 ± 0.16 mV and 1.02 ± 0.25 mV, respectively; *n* = 9 cells, paired t-test, *p* < 0.01). Right: Δ*V*_m_ variance summary for the teleportations within the familiar environment (FF) and the teleportations from the familiar to the novel environment (FN2) (−0.5 ± 0.4 mV^2^ and 1.4 ± 1.0 mV^2^, respectively; *n* = 9 cells, paired t-test, *p* > 0.05). **f** Left: the distribution of baseline *V*_m_ of granule cells recorded in control animals is represented as a green histogram (*n* = 73 cells). The right tail of a Gaussian fit for the dataset is shown in blue (Data fit). The orange curve shows the right tail of a Gaussian fit for a dataset depolarized by the experimentally observed mean value (1.0 mV) to estimate the effect of novelty on spiking (Novelty fit). The threshold above which 3.3% of the granule cell population produces spikes is indicated by the vertical red dashed line. Middle: enlarged view of the right tail of the Gaussian fits showing the area under the curve between the spiking threshold and 3 standard deviations from the mean. Right: diagram representing the percentage of spiking cells recruited from the silent population as calculated from the artificially depolarized dataset. See also Fig. S1.

### A small depolarization can lead to a large increase in the relative fraction of spiking cells

How does the observed synaptic response to novelty affect population activity in the dentate gyrus? To quantify this effect, we used our full dataset of baseline membrane potential values recorded in granule cells (*n* = 73 cells) in conjunction with recently published ground-truth data on dentate gyrus population activity^27^ to estimate that, within a 2-s time window, the fraction of spiking neurons increases from 3.3% to 4.2% (Fig. 2f; see Methods). This result indicates that a relatively small membrane potential depolarization observed at the individual neuron level, if driven by a generalized network mechanism, would entail the recruitment of a small but — in relative terms — sizeable amount of silent neurons to the active population. While the depolarization affects all granule cells equally, only those that are close to firing threshold will be recruited, providing specificity for cells that are synaptically activated upon transition to novelty. In addition, a small depolarization will also increase the firing rates of the small population of active neurons, and it may drive some granule cells into firing bursts of action potentials, as has previously been described *in vivo*^27,31^. Such burst firing would further amplify the effect of the transient depolarization on population activity.

### Isolated visual stimuli fail to depolarize granule cells

To probe if the observed synaptic response to saliency requires the animal’s engagement in the behavioral task, we presented isolated visual stimuli to untrained head-fixed mice moving freely on the treadmill surrounded by a uniformly dark screen (Fig. 3a). We then obtained whole-cell patch-clamp recordings from granule cells and presented periodical flashes of LED collimated light directed to the mice’s eyes (Fig. 3b). We computed the stimulus-aligned average within a 1 s window around the light flash presentation across multiple recordings and compared it to a bootstrap dataset (Fig. 3c,d). In this case, we did not observe a significant difference in membrane potential (Δ*V*_m_: 0.01 ± 0.10 mV for the data and 0.16 ± 0.08 mV for the bootstrap; *n* = 10 cells, paired t-test, *p* > 0.05; Fig. 3e) or in membrane potential variance (0.02 ± 0.21 mV^2^ for the data and −0.07 ± 0.18 mV^2^ for the bootstrap; *n* = 10 cells, paired t-test, *p* > 0.05; Fig. 3e) within this time window. This Δ*V*_m_ in response to light flashes was significantly different from the one observed during teleportations from the familiar to the novel environment (FN2) (FN2, *n* = 9 cells versus flashes, *n* = 10 cells; unpaired t-test, *p* < 0.01; Fig. 3f). The absence of a transient depolarization when isolated visual stimuli are presented suggests that saliency by itself is not sufficient to trigger the synaptic effect observed before. Instead, our finding indicates that the targeted locomotor engagement of the animal in the behavioral task is necessary for this form of synaptic novelty detection.

**Fig. 3.**
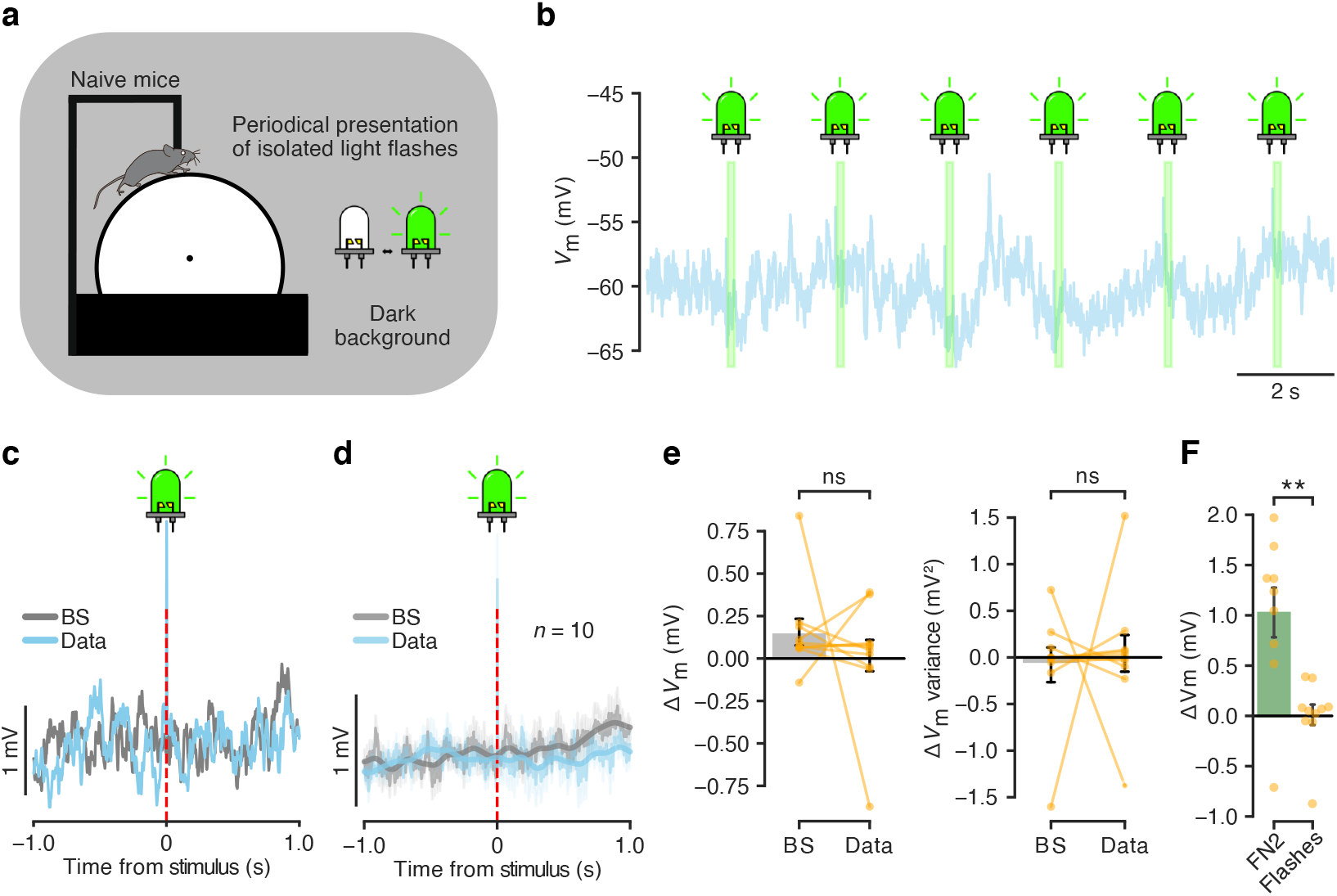
Isolated visual stimuli fail to depolarize granule cells. **a** Schematic of the experiment. Naive head-fixed mice move freely on a linear treadmill in a dark environment. Collimated LED light is flashed periodically on the eyes of the animals. **b** Example whole-cell recording from a granule cell during a light-flash experiment. The trace shows *V*_m_ and the light flash events are indicated in green. **c** Stimulus-aligned average from a representative recording. The blue trace represents the mean *V*_m_ aligned to the light flash events. The grey trace represents the mean *V*_m_ from a bootstrap dataset. The light flash event time is indicated by the vertical red dashed line (note also the presence of the stimulation artifact). **d** Stimulus-aligned average across multiple recordings. The blue trace represents the mean ± s.e.m. of *V*_m_ aligned to the light flash events. The grey trace represents the mean ± s.e.m. of *V*_m_ from a bootstrap dataset. The light flash event time is indicated by the vertical red dashed line (note also the presence of the stimulation artifact). Low-pass filtered traces (bold traces) are shown superimposed. **e** Left: Δ*V*_m_ summary for the bootstrap dataset events (BS) and the light flash events (Data) (0.16 ± 0.08 mV and 0.01 ± 0.10 mV, respectively; *n* = 10 cells, paired t-test, *p* > 0.05). Right: Δ*V*_m_ variance summary for the bootstrap dataset events (BS) and the light flash events (Data) (−0.07 ± 0.18 mV^2^ and 0.02 ± 0.21 mV^2^, respectively; *n* = 10 cells, paired t-test, *p* > 0.05). **f** Bar graph showing the Δ*V*_m_ summary for teleportations from the familiar to the novel environment (FN2) and in response to light flashes (Flashes) (FN2, *n* = 9 cells versus Flashes, *n* = 10 cells; unpaired t-test, *p* < 0.01). Left bar: same data as in Fig. 2e middle, right bar. Right bar: same data as in **e** left, right bar.

### Local blockade of muscarinic acetylcholine receptors abolishes the membrane potential response to saliency in granule cells

As the neuromodulatory transmitter acetylcholine is thought to play a key role in orchestrating the network changes related to attention and task engagement in response to uncertainty^32–34^, we repeated our experiment while blocking metabotropic acetylcholine receptors using the non-specific muscarinic competitive antagonist atropine through local stereotaxic injection immediately prior to the recordings. We confirmed the accuracy of our drug application technique and the extent of diffusion of the injected volume by injecting a solution containing a fluorescent dye (BODIPY) in a subset of animals (Fig. S2; see Methods). We did not observe a significant difference in the baseline intrinsic electrophysiological properties of the recorded cells after application of atropine when compared with our previous recordings (Baseline *V*_m_: atropine, −63 ± 2 mV, *n* = 8 cells; control, −68 ± 1 mV, *n* = 73 cells; unpaired t-test, *p* > 0.05. Baseline input resistance: atropine, 234 ± 16 MΩ, *n* = 8 cells; control, 216 ± 11 MΩ, *n* = 73 cells; unpaired t-test, *p* > 0.05. Fig. 4a). We then performed teleportation-aligned average analysis as described above (Fig. 4b-e). As observed in the experiment without drug application, we did not detect a significant difference between the mean membrane potential recorded during the total time spent in both virtual environments (−67.6 ± 2.2 mV in F and −67.5 ± 2.1 mV in N2; *n* = 6 cells, paired t-test, *p* > 0.05; Fig. 4e). Notably, we did not find a change in membrane potential or membrane potential variance between FF and FN2 teleportations when analyzing a 1 s time window after the teleportation events (Δ*V*_m_: −0.37 ± 0.19 mV for FN2 teleportations and −0.38 ± 0.36 mV for FF teleportations; *n* = 6 cells, paired t-test, *p* < 0.05. Δ*V*_m_ variance: −0.1 ± 0.7 mV^2^ for FN2 teleportations and 0.5 ± 0.5 mV^2^ for FF teleportations; *n* = 6 cells, paired t-test, *p* > 0.05; Fig. 4e). Moreover, the Δ*V*_m_ observed during FN2 teleportations after local injection of atropine was significantly different from the one observed in control animals (atropine: *n* = 6 cells; control: *n* = 9 cells; unpaired t-test, *p* < 0.01; Fig. 4f). The absence of membrane potential depolarization in response to novelty under muscarinic blockade suggests that this effect depends on metabotropic cholinergic signaling.

**Fig. 4.**
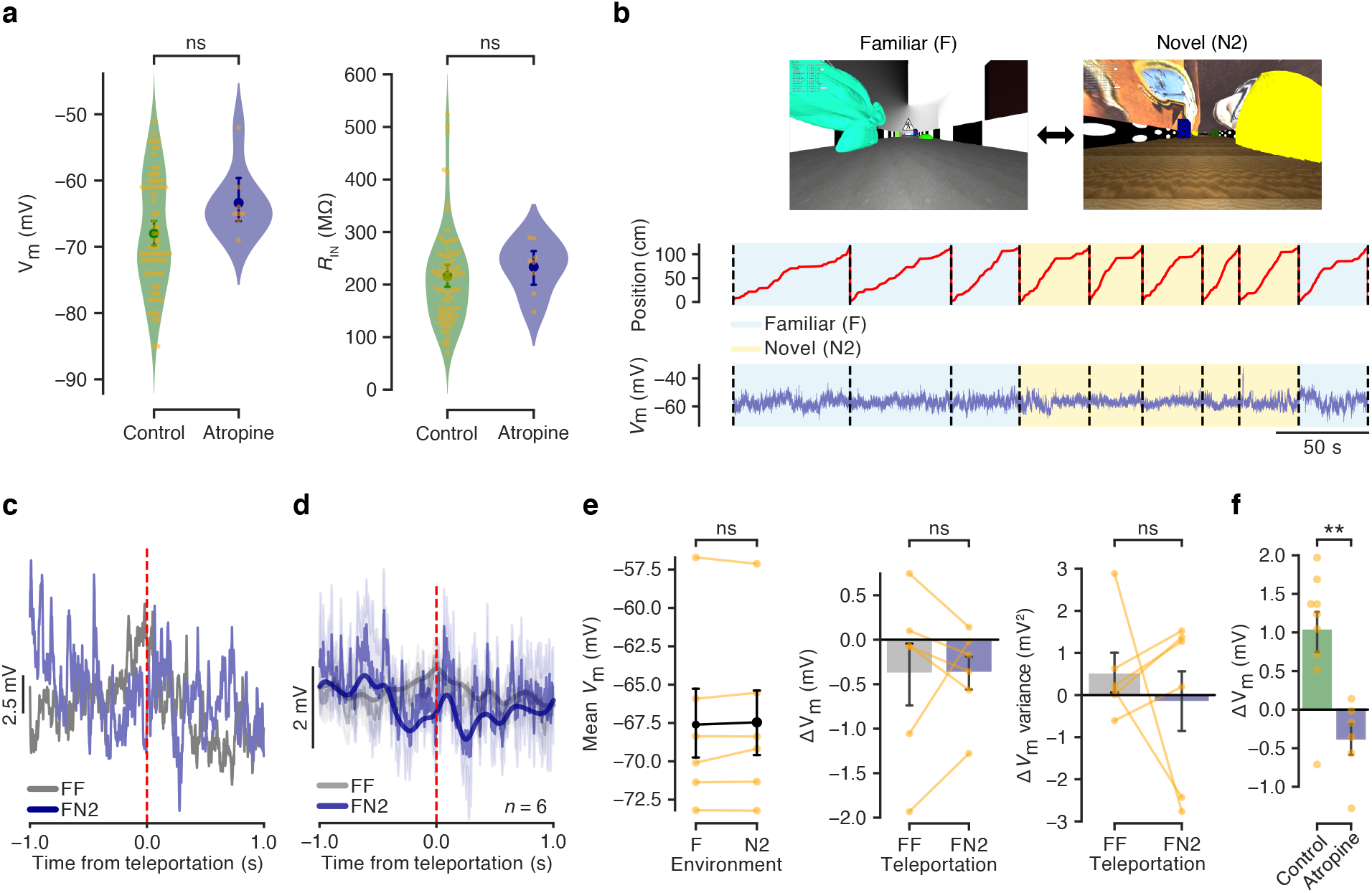
Local blockade of muscarinic acetylcholine receptors abolishes the membrane potential response to saliency in granule cells. **a** Left: baseline *V*_m_ summary for granule cells recorded in control animals and in animals after local injection of atropine (control: 68 ± 1 mV, *n* = 73 cells; atropine: 63 ± 2 mV, *n* = 8 cells; unpaired t-test, *p* > 0.05). Right: Summary of input resistances for granule cells recorded in control animals and in animals after local injection of atropine (control: 216 ± 11 MΩ, *n* = 73 cells; atropine: 234 ± 16 MΩ, *n* = 8; unpaired t-test, *p* > 0.05). **b** Example recording during a behavioral discrimination experiment switching between the familiar (F) and a novel environment (N2). Top: view along the long axis of the corridor in the familiar (F, left) and the novel 2 (N2, right) environments. Traces show animal position along the corridor (middle) and *V*_m_ (bottom). **c** Teleportation-aligned average from a representative recording. Traces represent the mean *V*_m_ aligned to the teleportation events. The average from teleportations within the familiar environment (FF) is shown in grey and the teleportation from the familiar to the novel environment (FN2) is shown in mauve. Teleportation time is indicated by the vertical red dashed line. **d** Teleportation-aligned average across multiple recordings in animals after local injection of atropine. Traces represent the mean ± s.e.m. of the *V*_m_ aligned to the teleportation events. Teleportations within the familiar environment (FF) are shown in grey and teleportations from the familiar to the novel environment (FN2) are shown in mauve. The teleportation event time is indicated by the vertical red dashed line. Low-pass filtered traces (bold traces) are shown superimposed. **e** Left: Mean *V*_m_ summary for the familiar and the novel (N2) environments after local injection of atropine (−67.6 ± 2.2 mV and −67.5 ± 2.1 mV, respectively; *n* = 6 cells, paired t-test, *p* > 0.05). Middle: Δ*V*_m_ summary for the teleportations within the familiar environment (FF) and the teleportations from the familiar to the novel environment (FN2) after local injection of atropine (−0.38 ± 0.36 mV and −0.37 ± 0.19 mV, respectively; *n* = 6 cells, paired t-test, *p* > 0.05). Right: Δ*V*_m_ variance summary for the teleportations within the familiar environment (FF) and the teleportations from the familiar to the novel environment (FN2) after local injection of atropine (0.5 ± 0.5 mV^2^ and −0.1 ± 0.7 mV^2^, respectively; *n* = 6 cells, paired t-test, *p* > 0.05). **f** Bar graph showing the Δ*V*_m_ summary for teleportations from the familiar to the novel environment (FN2) in control animals and in animals after local injection of atropine (Control, *n* = 9 cells versus Atropine, *n* = 6 cells; unpaired t-test, *p* < 0.01). Left bar: same data as in Fig. 2e middle, right bar. Right bar: same data as in **e** middle, right bar. See also Fig. S2.

### A computational model suggests that increased dentate gyrus activity can initiate a map switch in CA3

To investigate the potential downstream effects of the transient increase in granule cell activity, we implemented a network model of CA3, subject to inputs from the dentate gyrus and the medial entorhinal cortex (mEC). The model is based on the continuous attractor theory of spatial memory and navigation^35^. Place cells in the CA3 model network are supported by excitatory inputs from mEC, in correspondence with the physical position of the rodent in a specific environment, and by the recurrent connections that match the statistics of these inputs. The learning process producing these connections through repeated exposure to the same environment is not explicitly modeled here. The pattern of these connections forms a single consolidated map of the familiar environment (F), as well as a large, weakly and irregularly pre-wired subnetwork that could provide representations of a novel one (N)^36^ (Fig. 5a; see Methods). Random projections from the dentate gyrus to CA3 neurons transiently excite a fraction of map N neurons following ‘teleportation’ and contain no spatial selectivity (Fig. 5b). Furthermore, in our model the dentate gyrus recruits feed-forward inhibition in a frequency-dependent manner^37,38^.

**Fig. 5.**
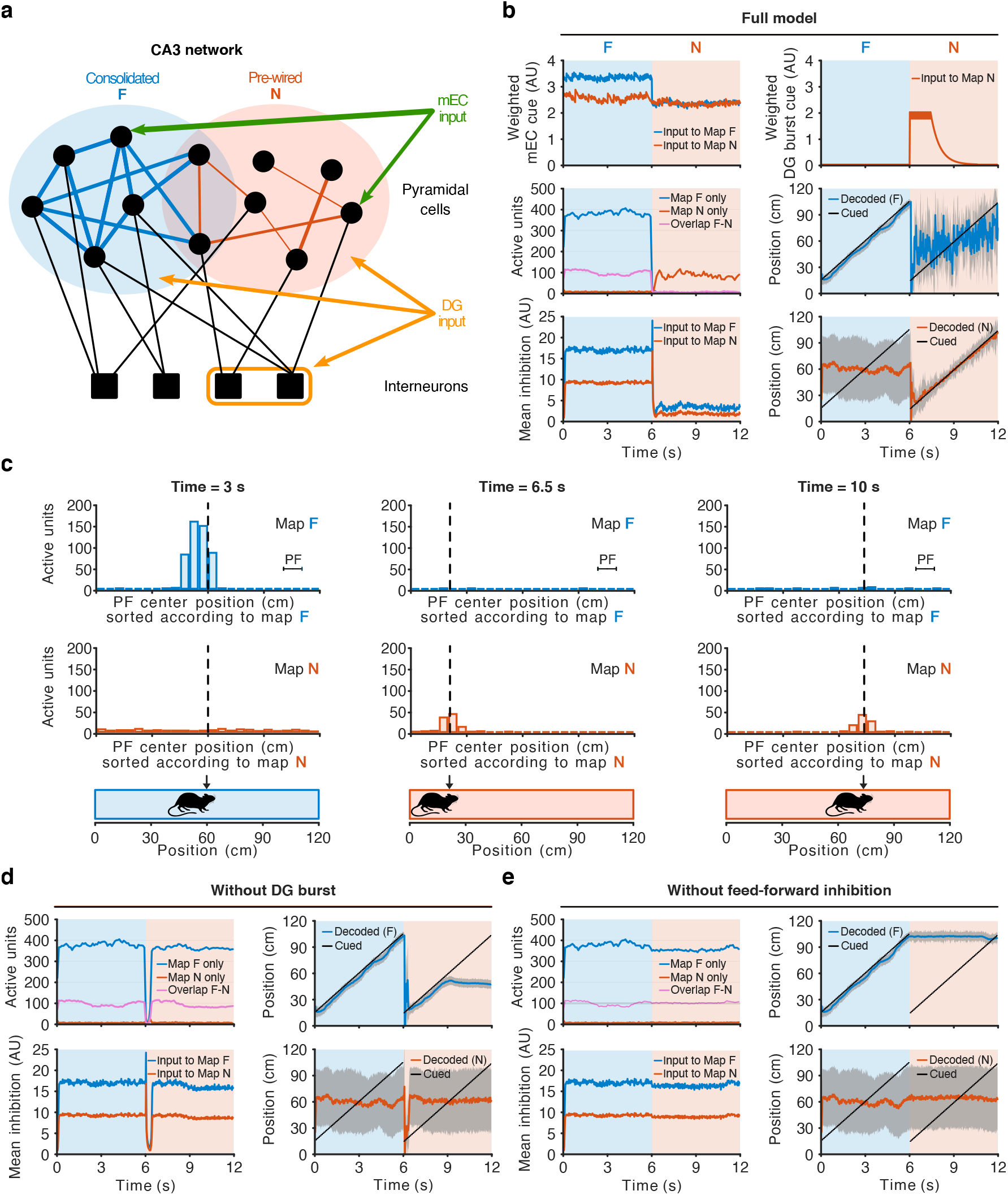
A computational model suggests that increased dentate gyrus activity can initiate a map switch in CA3. **a** CA3 network model. Consolidated (blue) and irregular, weak (red) recurrent excitatory connections between place cells (black dots) support, respectively, the consolidated (familiar F, blue) and pre-wired (novel N, red) maps. Place cells receive feed-forward (from DG) and recurrent inhibitory inputs from interneurons (black squares), and excitatory inputs from mEC (green) and DG (orange). **b** Top row: Dynamics of external inputs to the CA3 cells. Excitatory inputs are subdivided into spatially-related mEC (left) and transient DG (right) inputs. Middle and bottom rows: Network dynamics. (Middle left) Number of active cells in CA3 plotted against time. Neurons with a place field in only one of the two maps are plotted in blue (F map) or red (N map). Active units with a place field in both maps are represented in purple. (Bottom left) Inhibitory inputs result from the combined effects of global and novelty-driven inhibition. (Right: middle and bottom): comparison between the cued position (black) and the one decoded from the active neurons in maps F (blue, middle) and N (red, bottom). Decoded positions were calculated by averaging the position of the place field centers of active cells in the two maps. Grey shaded areas represent the standard deviation associated with the decoded position. **c** Snapshots of the activity for three animal positions (bottom row) in F (left), right after teleportation (center), and in N (right). Blue and red bars represent the number of active cells with a place field center in the corresponding 5 cm bin in, respectively, the F (first row) and N (second row) maps. Black dashed lines represent the position cued through the mEC input to the subnetworks involved in the encoding of the two maps. **d-e** Network dynamics for incomplete models (similar to panel **b** middle and bottom rows). **d** Model without the novelty-driven excitatory input from DG. When the transient inhibition accompanying teleportation to novelty ends, a stable bump re-emerges in map F, due to the stronger recurrent connections. **e** Model without the novelty-activated feed-forward inhibition from DG. The activity bump in the F map remains in its last position before the teleportation and prevents the formation of a new bump in map N. In both cases lower mEC inputs to map F in the novel environment (panel **b**, top left) are not sufficient to consistently follow the animal position. See also Figs. S3 and S4 and Supplementary Video.

In the familiar (F) environment, place cells are activated by mEC inputs and the strong recurrent connections in the network supporting map F. This connectivity results in a large bump of activity, coding at all times for the position of the animal as it moves along the track (see Fig. 5b, middle right and 5c, left). Note that even if the rodent is in the familiar environment, mEC inputs excite some place cells which are shared with the immature ‘pre-map’ N (purple line in Fig. 5b, middle left), but are not sufficient to trigger another activity bump there (Fig. 5b, bottom right), as the global inhibition of the CA3 network allows for only one activity bump at a time. Following teleportation to the novel environment, two phenomena take place as a result of the burst of activity in the dentate gyrus. On the one hand, activity in the CA3 network is temporarily strongly inhibited by feedforward inputs^39^ (Fig. 5b, bottom left), abolishing the activity bump in map F (Fig. 5c, middle). On the other hand, the enhanced inputs from dentate gyrus towards pre-map N (Fig. 5b, top right) randomly stimulate additional cells in pre-map N. This boost in activity initiates a coherent, self-sustained new bump of activity around the neurons receiving most inputs (Fig. 5c, middle). This new neural activity bump then persists after dentate gyrus inputs return to their baseline value, and codes robustly for the animal’s position in the novel map at all times (Fig. 5c, right), even though the inputs from mEC in the novel map are not stronger than the ones pointing towards the familiar map (Fig. 5b, top left).

These two-fold effects of the increase in activity in the dentate gyrus are crucial for the coherent switch from map F to pre-map N to take place (see Supplementary Video). If no burst of excitatory inputs from the dentate gyrus to pre-map N accompanies the transition, the bump in map F is reinstated after its transient disappearance as the animal navigates in the novel environment (Fig. 5d). Furthermore, in the absence of transient inhibition in CA3, the bump of activity in map F is not erased and persists in the novel environment (Fig. 5e). Thus, the model shows that transient inputs from the dentate gyrus are essential to switch from a robust familiar representation to a weakly pre-formed novel map in the downstream attractor network.

## Discussion

Here we show that dentate gyrus granule cells exhibit a transient small depolarization of their membrane potential when animals are ‘teleported’ into a novel virtual environment. By contrast, isolated visual stimuli that are presented independently of the animal’s behavior fail to evoke any systematic synaptic response. The observed novelty-dependent depolarization can be blocked by local application of atropine, indicating that it depends on activation of muscarinic (metabotropic) acetylcholine receptors. While the amplitude of the depolarization is only on the order of 1 mV, because of the sparse activity levels in the dentate gyrus even such a small depolarization can have a large effect on the relative increase in the proportion of firing neurons. Computational modeling reveals that this increased activity level in the dentate gyrus can push the downstream CA3 network from retrieving an existing representation of a familiar environment to forming a new one of the novel environment.

To support ongoing behavior in an environment with mixed novel and familiar components, hippocampal circuits need to be able to store relevant novel information without disrupting the recall of familiar memories^21–23^. It has been suggested that upon transition into a novel environment, robustly decorrelated inputs from the dentate gyrus push downstream circuits to form a new neuronal representation (i.e. ‘global remapping’). However, the attractor dynamics of the CA3 circuit counteract this process by stabilizing the familiar representation as the default mode^2,8,20,21^, thereby putting the two processes at odds with each other. Additional inputs containing information about the saliency and the unexpectedness of the novel environment^32^ may provide the required arbiter signal that decides whether the CA3 circuit forms a new neuronal representation or recalls a stored memory.

How is this ‘novelty signal’ conveyed to the hippocampal network? In contrast to the multimodal cortical inputs to the hippocampus, which carry information about distinct memory elements, subcortical inputs are known to contain information about ‘global’ internal brain states, such as attention, uncertainty and arousal^24,25,32^. Signals of this sort could be transmitted to the hippocampus in the form of a generalized short-lived network perturbation that could in turn alter the manner in which information is processed in the circuit. This kind of global transitory change in circuit dynamics can occur in response to diverse neuromodulatory transmitters, which are crucial evolutionary-conserved elements for the reshaping and repurposing of neural circuits during associative and non-associative learning^25,40,41^. Different neuromodulatory systems appear to play distinct roles during mnemonic processing and, among these, cholinergic neuromodulation has been suggested to be the one in charge of signaling ‘absolute’ novelty to the hippocampus^22,42^. Indeed, theoretical models have suggested that increased cholinergic neuromodulation promotes encoding of novel information in the hippocampal network while reduced cholinergic neuromodulation favors consolidation of previously encoded patterns^33,42,43^. This notion is supported by experimental evidence of a rise in acetylcholine levels in the hippocampus when an animal encounters a novel spatial environment^44–47^ and of memory impairment by pharmacological blockade of cholinergic neurotransmission^48–50^, which selectively affects encoding while sparing retrieval^51^.

Previous studies have shown that optogenetic stimulation of cholinergic septohippocampal projections can recruit excitatory inputs to granule cells as a component of a bimodal synaptic response^52^. The inhibitory late component of this response is mediated by multiple intermediaries, including astrocytes and local interneurons, depends on nicotinic acetylcholine and GABA_A_ receptor channels, and can be revealed at a negative chloride reversal potential (~ −88 mV) when septohippocampal fibers are synchronously stimulated at high frequencies. In our recordings, which were obtained at a physiological *E*_Cl_ for mature granule cells (~ −72 mV)^53,54^, the excitatory component of the response appears to be isolated, as hyperpolarizing inhibition may be absent due to the small driving force for chloride ions, low-frequency asynchronous activation of cholinergic inputs *in vivo*, and differences in synaptic transmission dynamics, such as baseline GABAergic tone, in awake versus anesthetized animals^55,56^. Thus, while the transient short-lived signal that we observe is depolarizing, potential shunting inhibition appearing at a later phase, depending on glial intermediaries, may explain why an increased cholinergic tone can lead to reduced overall granule cell activity in novel environments under some conditions^10,57^.

The depolarization that we observe is sensitive to local application of atropine, suggesting that it depends on metabotropic acetylcholine receptors, consistent with the temporal dynamics of the signal. From the five subtypes of muscarinic receptors, M1, M2 and M4 are the most widely expressed in the hippocampus^58^. Among these, the M2 subtype is expressed only in interneurons. On the other hand, the M1 subtype is preferentially expressed in the somatodendritic compartment of principal cells^59,60^ and is known to enhance postsynaptic excitability and NMDA receptor activity by inhibiting potassium channels^61,62^ and calcium-activated SK channels^63,64^. These features make it a candidate driver of the membrane potential depolarization that we observe in granule cells^34,48,50^, in accordance with the disappearance of the effect observed in our experiments when these receptors are blocked with atropine. Nonetheless, it has recently been reported that hilar mossy cells — which are also known to be targets of cholinergic neuromodulation^65,66^ — play a key role during novelty detection in the hippocampus^67^, which raises the possibility that the synaptic response to novelty that we recorded in granule cells is not driven by the direct effect of acetylcholine on granule cells but instead indirectly through cholinergic modulation of local interactions in the intrahippocampal circuit. Genetically targeted manipulation of specific nicotinic and muscarinic acetylcholine receptors in combination with cell-specific neuronal recordings during novelty-associated behavior will be necessary to clarify the precise mechanisms whereby acetylcholine modifies the synaptic properties of the hippocampal circuit in response to saliency.

How does the synaptic novelty signal affect the activity of individual granule cells? The small depolarization that we observe will selectively affect the firing of neurons that are already synaptically activated: it will increase the firing rate of neurons that are actively spiking, and it will drive silent cells to fire spikes if their membrane potential is close to the action potential threshold. It has been shown that evoking spikes in previously silent granule cells *in vivo* at specific locations in the environment can lead to the induction of place fields, especially under novelty^68^. We therefore expect that at least part of the newly recruited active neurons will permanently fire upon transition into the environment that led to their activation when the animal first encountered it, even as the environment grows familiar. Such a process may facilitate the robust retrieval of the corresponding attractor state in CA3 over time.

Studies on the activity of granule cell populations have reported both decreased and increased activity as an animal familiarizes itself with a novel environment over the time course of several minutes and more, and the long-term dynamics of this process remains to be delineated^10,57,67^. However, how the population activity changes at the moment when the animal transitions into a novel environment is unclear. As the overall activity levels in the dentate gyrus are notably sparse, the transient depolarization will only recruit a small absolute number of neurons into the spiking population, and may only lead to firing of few or even single additional action potentials. Population recording techniques that are typically employed in virtual reality, such as 2-photon imaging^10,12^, are likely to miss these additional few spikes that occur only in a small number of neurons during a short time window^69^. Highly sensitive electrophysiological recordings with high temporal and single-spike resolution from large populations of neurons during instantaneous environment switches in virtual reality will be required to observe the predicted population response^70^.

How can a transient increase in dentate gyrus activity affect downstream circuits? By implementing a computational model of CA3, we reveal that a small and short excitation from the dentate gyrus, accompanied by transient increased inhibition^39^, can initiate a weak but self-sustained activity bump that encodes the rodent position in a non-consolidated, pre-configured map of the novel environment. Both the transient fast inhibition and the slower excitation required in this model can result from increased activity in the dentate gyrus, as mossy fibers provide both monosynaptic excitation as well as disynaptic feedforward inhibition to CA3 pyramidal cells. The excitation-inhibition balance is governed by complex frequency-dependent dynamics, with net inhibition predominating at low presynaptic firing frequencies and a switch to net excitation occurring as frequency increases^37,38^. Thus, the ability to invert the polarity of the mossy fiber-CA3 pyramidal neuron synapse in a frequency-dependent manner may be necessary when switching between information processing modes in the hippocampal circuit under the regulation of extrahippocampal signals. Moreover, mossy fiber-CA3 synapses show pronounced post-tetanic potentiation in response to natural bursting activity patterns described *in vivo*^27,31,71^, which could be triggered by the small depolarization that we observe. This form of synaptic plasticity would further boost excitatory drive, supporting the new activity bump.

The existence of pre-configured assemblies is at the basis of the so-called preplay phenomenon^72^, and is compatible with the observed emergence of alternative maps under silencing of place-cell assemblies^36^. Contrary to previous models of switching between two equally consolidated maps^73^, switching to an immature map crucially requires transient inhibition, in agreement with reports that somatic inhibition is transiently increased following novelty in CA1^39,74^. Learning processes, possibly involving reconfiguration of inhibitory circuits^75^, could ultimately strengthen and reshape this primitive, pre-configured representation.

In the model that we propose, cholinergic modulation of the dentate gyrus affects network dynamics downstream in area CA3. What could be the benefit of separating the location of the modulation and its effect, instead of directly activating CA3 neurons? One advantage is that only those granule cells that are synaptically activated during transition to novelty, bringing them close to or above the firing threshold, are specifically modulated. As the dentate gyrus produces codes that strongly discriminate between familiar and novel environments, only a subset of inputs to CA3 that is highly specific for the novel environment increases its activity. A novelty signal that non-specifically acted on CA3 neurons would activate, among others, assemblies coding for the familiar environment, as their activity is maintained by their attractor properties, even if the animal is already in a novel environment. This stable attractor state would then prevent novel representations from forming in CA3. Another advantage is that a transient increase in the fraction of active neurons enhances the separability of representations^76^. The depolarization that we observe in the dentate gyrus in response to novelty could thus temporarily increase the discriminative power of the network, further favoring the establishment of a new neuronal assembly in CA3. Furthermore, activating inputs from the dentate gyrus to CA3 that are active or close to spike threshold during teleportation to novel environments may induce Hebbian plasticity at specific DG-to-CA3 synapses, so that novel information is rapidly stored in specific assemblies. The upstream cholinergic processes in the dentate gyrus that we describe here are not in conflict with previously reported cholinergic modulation of CA3^1^; on the contrary, we suggest that they synergistically promote plasticity.

Neuronal generalization and discrimination are processes that must occur in concert, as familiar memories need to be robustly retrieved but also updated with relevant novel information. In addition, any experience consists of multiple elements in different physical dimensions that may need to be stored as separate memories depending on their behavioral relevance^21–23^. Inducing a small and transient universal ‘bias’ in the population code of the dentate gyrus when faced with novelty may provide a solution to this challenge, as the overall structure of the dentate gyrus population code is only weakly and briefly affected while the novel environment is learnt and familiarized. Novelty can thereby flexibly tag different dimensions of an experience to produce multifaceted memory representations (Fig. 6).

**Fig. 6.**
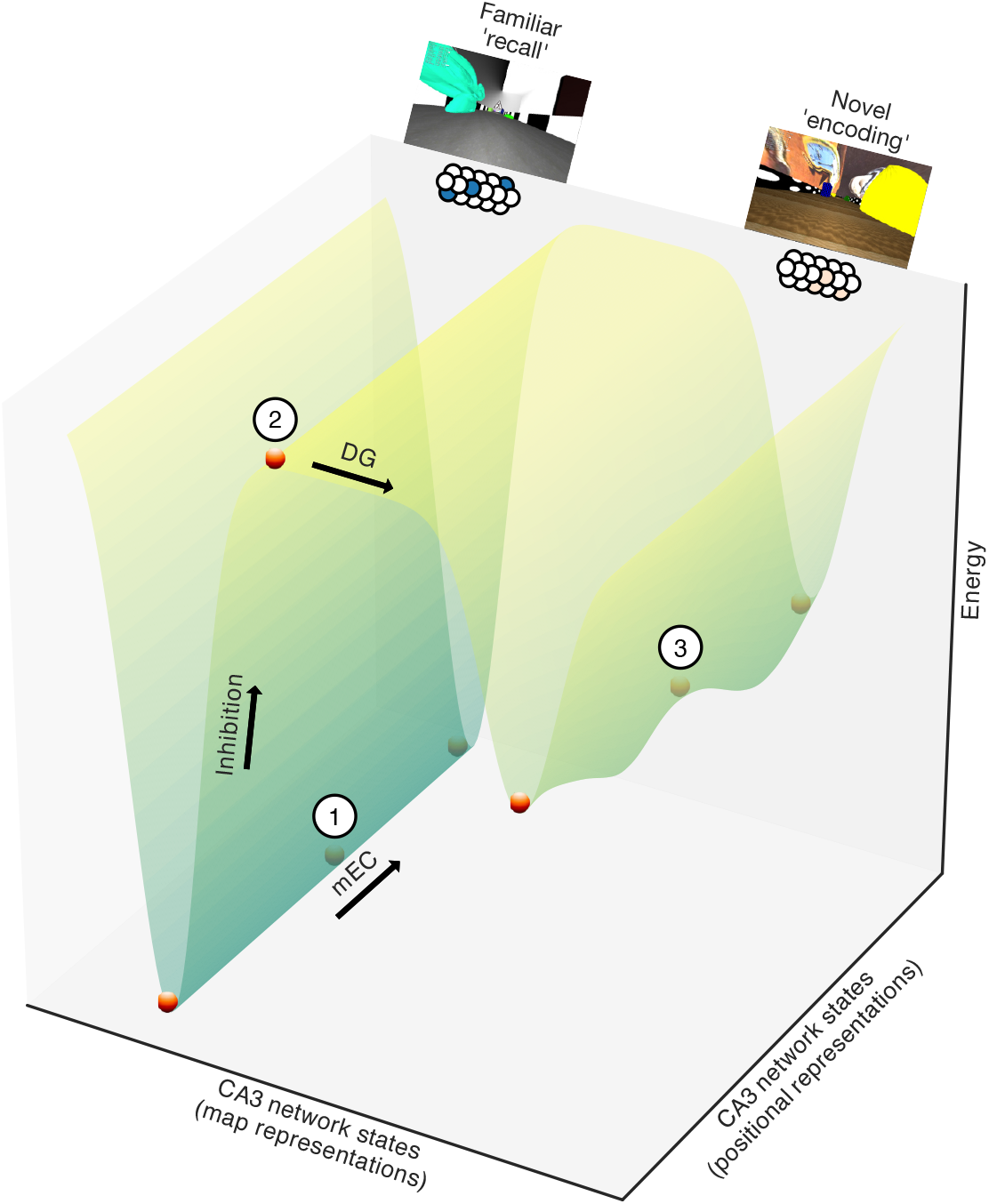
Switching between familiar and novel attractor states: proposed model. Energy landscape of CA3 network states representing different maps and different positions along the track (schematic). In a familiar environment, the CA3 network falls into an attractor state that is governed by strengthened recurrent synaptic connectivity, thereby performing generalization during memory recall of familiar events. We propose that a small bias in the inputs from the dentate gyrus first recruits feed-forward inhibition, thereby lifting the network state out of the deep trough representing the familiar environment. Direct excitation from the dentate gyrus then pushes the CA3 network into a different attractor state with initially weaker, pre-existing recurrent connectivity, thereby performing discrimination during novelty encoding. Numbers represent network states corresponding to the model snapshots shown in Fig. 5c (1: Fig. 5c, left; 2: Fig. 5c, middle; 3: Fig. 5c, right).

## Methods

### Animals and surgical procedures

All procedures were conducted in accordance with European and French regulations on the ethical use of laboratory animals for experimentation (EU Directive 2010/63/EU) and were reviewed and approved by the Ethics Committee of the Institut Pasteur CETEA (project number dap160066). 5-16 week-old male C57BL/6J wild-type mice (Janvier Labs) were used. Animals were housed in groups of four in polycarbonate individually-ventilated cages equipped with running wheels and were kept under a 12-h inverted light/dark cycle with *ad libitum* access to food and water. All animals were treated identically. Multimodal analgesia (buprenorphine 0.05 mg/kg + meloxicam 5 mg/kg) was administered through intraperitoneal injection at least 30 minutes before any surgical intervention and the skin on the surgical area was infiltrated with lidocaine prior to incision. Antisepsis of the incision sites was performed with povidone-iodine (Betadine). Animals were anesthetized with isoflurane (3% for induction, 1-2% for maintenance, vol/vol) and placed in a stereotaxic apparatus (Kopf Instruments). The corneal surfaces were protected from desiccation by applying artificial tear ointment and the body temperature was kept constant throughout the surgical intervention with a heating pad. The skin was incised with scissors and the periosteum was mechanically removed using a surgical bone scraper. Stainless-steel headposts (Luigs & Neumann) were attached to the animals’ skulls using dental acrylic (Super-Bond C&B, Sun Medical). Postoperative analgesia (meloxicam 5 mg/kg) was administered orally in combination with surgical recovery feeding gel (ClearH_2_O). Animals were allowed to recover from surgery for at least seven days preceding the start of the training sessions.

### Behavioral tasks in virtual reality

Three custom virtual reality environments were developed using the Blender Game Engine (http://www.blender.org) in conjunction with the Blender Python Application Programming Interface. All environments consisted of a 1.2 m-long linear corridor visually enriched with proximal and distal cues and floor and wall textures. The reward delivery trigger zone was placed in an un-cued location of the corridor that was identical in all environments. The warped environments were projected onto a spherical dome screen (*ϕ* 120 cm) using a quarter-sphere mirror (*ϕ* 45 cm) placed underneath the mouse, as described previously . The screen covered ~240°, which corresponds to nearly the entire horizontal field of view of the mouse. Animals were head-fixed and placed on an air-supported polystyrene rolling cylinder (*ϕ* 20 cm) that they used as a treadmill to navigate the virtual scene. Cylinder rotation associated with animal locomotion was read out with a computer mouse (Logitech G500) and linearly converted to one-dimensional movement along the virtual reality corridor. Animals were extensively handled and habituated to the virtual reality setup before the onset of experimental procedures. Animals underwent 5 training sessions of 20-30 min each on consecutive days prior to the electrophysiological recordings. All training sessions were conducted during the dark phase of the light cycle of the mice. During the training period and the experiments, animals were water-restricted to 80% of their baseline weight to maximize their behavioral drive^80^. Body weight and general health status were monitored daily. Animals were trained to navigate the virtual corridor in the familiar environment (F) and to retrieve an 8% sucrose solution reward by stopping for at least 3 s in an un-cued reward zone placed at a fixed location of the corridor. Licking behavior was monitored using a custom-made Arduino piezoelectric sensor coupled to the reward delivery spout. Animals were ‘teleported’ back to the beginning of the track upon crossing of a defined threshold near the end of the virtual corridor. As the virtual environments, training protocol and reward contingencies used in this study are different from the ones used in previous work^12^, the hit rate results are not directly comparable. After having completed the five-day training protocol in the familiar environment, a behavioral recording session was conducted in which laps in the familiar environment (F) were alternated with laps in the novel environment 1 (N1). The same environment alternation strategy was used during the electrophysiological recordings, using the familiar environment (F) and the novel environment 2 (N2). A purely behavioral session was conducted separately from the electrophysiological recordings since the latter are typically very short and therefore do not provide enough behavioral data to accomplish an accurate assessment of task performance.

### *In vivo* whole-cell patch-clamp recordings

Two craniotomies (*ϕ* ~0.5 mm) were drilled 3-24 hours before the recording session for the recording electrode (right hemisphere, 2.0 mm caudal and 1.5 mm lateral from Bregma) and the reference electrode (left hemisphere, 2.0 mm caudal and −1.5 mm lateral from Bregma). The *dura mater* was removed using fine forceps and the cortical surface was kept covered with artificial cerebrospinal fluid of the following composition: 150 mmol/L NaCl, 2.5 mmol/L KCl, 10 mmol/L HEPES, 2 mmol/L CaCl_2_, 1 mmol/L MgCl_2_. In a subset of animals, 600 nL of 1 mmol/L atropine solution was injected with a glass micropipette at a depth of 1.7 mm from the cortical surface to selectively target the dentate gyrus^81^. Recording electrodes were pulled from filamented borosilicate glass (Sutter Instrument) and filled with internal solution of the following composition: 135 mmol/L potassium methanesulfonate, 7 mmol/L KCl, 0.3 mmol/L MgCl_2_, 10 mmol/L HEPES, 0.1 mmol/L EGTA, 3.0 mmol/L Na_2_ATP, 0.3 mmol/L NaGTP, 1 mmol/L sodium phosphocreatine and 5 mg/mL biocytin, with pH adjusted to 7.2 with KOH. All chemicals were purchased from Sigma. Pipette tip resistance was 4-8 MΩ. Electrodes were arranged to penetrate the brain tissue perpendicularly to the cortical surface at the center of the craniotomy and the depth of the recorded cell was estimated from the distance advanced with the micromanipulator (Luigs & Neuman), taking as a reference the point where the recording electrode made contact with the cortical surface. Whole-cell patch-clamp recordings were obtained using a standard blind-patch approach, as previously described^12,27,82^. Only recordings with a seal resistance > 1 GΩ were included in the analysis. Recordings were obtained in current-clamp mode with no holding current. No correction was applied for the liquid junction potential. Typical recording durations were ~5 min, although longer recordings (~30 min) were occasionally obtained. *V*_m_ signals were low-pass filtered at 10 kHz and acquired at 50 kHz. After completion of a recording, the patch recording electrode was gently withdrawn to obtain an outside-out patch to verify the integrity of the seal and ensure the quality of the biocytin filling. To synchronize behavioral and electrophysiological recordings, TTL pulses were triggered by the virtual reality system whenever a new frame was displayed (frame rate: 100 Hz) and recorded with both the behavioral and the electrophysiological acquisition systems.

### Histology and microscopy

Immediately upon completion of a successful recording, animals were deeply anesthetized with an overdose of ketamine/xylazine administered intraperitoneally and promptly perfused transcardially with 1x phosphate-buffered saline followed by 4% paraformaldehyde solution. Brains were extracted and kept immersed overnight in 4% paraformaldehyde solution. 60-70 μm-thick coronal slices were prepared from the recorded hippocampi. Slices were stained with Alexa Fluor 488–streptavidin to reveal biocytin-filled neurons and patch electrode tracts. DAPI was applied as a nuclear stain to reveal the general anatomy of the preparation. Fluorescence images were acquired using a spinning-disc confocal microscope (Opterra, Bruker) and analyzed using ImageJ. The accuracy of the recording coordinates was confirmed in all cases by identification of either the recorded neuron or the recording electrode tract.

### Estimation of the effect of the observed synaptic response to novelty on population activity in the dentate gyrus

A Gaussian fit was produced for the complete dataset of baseline values of membrane potential recorded in granule cells (*n* = 73 cells). Then, an artificial dataset representing ‘novelty’ was generated by applying the observed mean depolarization (1.0 mV) to these baseline values and the corresponding Gaussian fit was produced. Ground-truth data on the activity of granule cells during spatial navigation was used to estimate that, during navigation in a familiar environment, ~3.3% of the granule cell population produces spikes during a 2 s period^27^. The right tail of the Gaussian fit from the real dataset was used to calculate the spiking threshold that would yield this percentage (−44.0 mV). Using this threshold, the percentage of cells that would be above it (i.e. actively spiking) in the artificially depolarized dataset was computed, which yielded 4.2%, representing a recruitment of spiking cells of approximately a third of the baseline spiking population’s size.

### CA3 network model: connections and inputs

An auto-associative neural network of *n* = 20,000 binary (0,1) units was implemented as a model of CA3. 20% of the *n* units were randomly assigned placed fields (PF) in environment F, uniformly centered on *p* = 400 equidistant points along the track (PF diameter = 33 pts). A scale factor of 0.3 cm was multiplied with this spatial dimension to simulate the physical length of the track (120 cm) in the experiment (PF diameter = 9.9 cm). Activity patterns *ξ^μ^* were generated for each point by randomly choosing 330 units (sparsity level *a* = 0.0165)^9^ among those with a PF overlapping the position. Parameters were adapted from Guzman et al. for a smaller number of total units by keeping fixed the absolute number of units involved in a single memory.

The coupling matrix *J* was defined through the clipped Hebb rule,

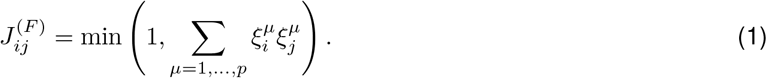

Such couplings carve a quasi-continuous attractor model of the environment^35^. Couplings 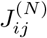 supporting the pre-wired map N were defined in the same way, based on another random subset of 0.2 *n* place cells, and were multiplicatively shrunk by random factors < 1 (beta-distribution, parameters *α* = 0.7, *β* = 1.2). As a result, all connections 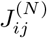 were reduced in strength, or even set to zero (Fig. S3). Finally, the excitatory synaptic matrix *J* for the CA3 network was defined as

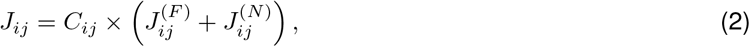

where the connectivity matrix *C_ij_* = 0,1 randomly assigned 1,200 input connections *j* to each unit *i*, in agreement with estimates of the connectivity^9^. mEC inputs to CA3 were spatially selective, acting on 50% of the place cells, chosen at random, among those involved in the activity pattern *ξ^μ^* associated with the current position of a virtual rodent. To account for consolidation of map F, input intensities on cells coding for environment F were stronger than for environment N while the rodent was navigating environment F. Conversely, while navigating environment N, mEC inputs of equal strengths were sent to the two maps. Dentate gyrus inputs, activated by novelty, acted on randomly chosen 2% of units in the subnetwork supporting pre-map N, with the same intensity as mEC inputs in the novel environment. Dentate gyrus baseline activity instead cued a random 2% fraction of CA3 units at all times, with no spatial or map selectivity.

### CA3 network dynamics

Place cells activities were updated by comparing the sum of their inputs to an activation threshold *G* = 2.31^9^ according to the following probability:

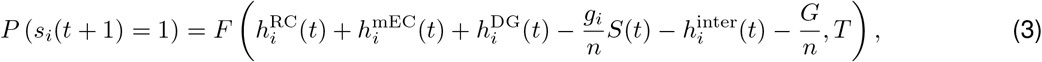

where *i* is the index of the unit, *s_i_* = 0,1 its activity, and *F* is the sigmoidal function 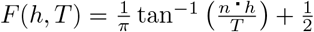,with the temperature *T* = 0.1, in agreement with the order of magnitude estimated for similar network models^83^. In addition to inputs 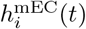 from mEC and 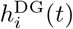 from dentate gyrus, cells received inputs through CA3 recurrent couplings, 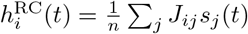, a global inhibition component proportional to network activity *S*(*t*), with *g_i_* = 0.035 for all units contributing to the memory of the F map and *g_i_* = 0.015 for units involved only in the storage of the N map, and inputs from *n_I_* = 5000 interneurons, 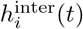. To balance the DG baseline input to CA3, global inhibition was set to have a minimal value counterbalancing the activity of *S*_min_ = 50 neurons, i.e. *S*(*t*) = max (*S*_min_, Σ_*j*_ *s_j_*(*t*)). Interneurons were modeled as threshold-linear units (threshold *G_I_* = 0.5), and sent inhibitory inputs to 500 place cells each; they were activated during teleportation to environment N through an external stimulation from the DG, effectively modeling a mechanism of feed-forward inhibition to CA3 linked to the transient increase of DG activity. The intensities of all external cues (i.e., mEC, DG, inter) decayed exponentially in time after each update of the corresponding cues, namely

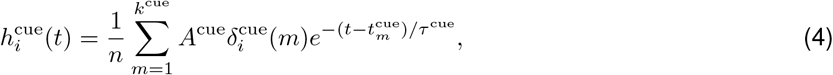

where *k*^cue^ is the number of times the intensity is refreshed after an update of the cue, 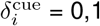 indicates if unit *i* receives the input, *A*^cue^ is the initial intensity, 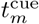 is the time of the *m*^th^ refresh for each update of the cue, and *τ*^cue^ is the decay time. In our simulations, we used *k*^mEC^ = *k*^inter^ = 1, *k*^DG^ = 15; *A*^mEC^ = 3 for map F during navigation of environment F (*A*^mEC^ = 2 while in environment N) and *A*^mEC^ = 2 for pre-map N (for navigation in both environments), *A*^DG^ = 2, *A*^inter^ = 20; *τ* ^mEC^ = 100, *τ* ^DG^ = 30, *τ* ^inter^ = 3. mEC and baseline DG cues were updated once every five time steps, DG burst activity was updated 15 times after the teleportation event (once every 5 time steps) and inter was updated only once at the teleportation time. Alternative sets of parameters were also tested to check the stability of the model (Fig. S4). In figures, time steps were scaled by a factor of 20 ms to match the average speed of the rodent along the track.

### Data analysis and statistics

To analyze subthreshold membrane potential, *V*_m_ traces were digitally low-pass filtered at 5 kHz and resampled at 10 kHz. *V*_m_ traces were subsequently high-pass filtered at > 10^−5^ Hz to remove slow trends such as reference drifts. Action potentials were removed from the traces by thresholding to determine action potential times and then replacing 2 ms before and 10-20 ms (depending on the action potential shape) after the action potential peak with an interpolated straight line. Data are presented as the mean ± s.e.m., unless stated otherwise. Statistical significance was assessed using either paired or unpaired two-tailed Student’s t-tests, as appropriate. Indications of statistical significance correspond to the following values: ns *p* > 0.05, * *p* < 0.05, ** *p* < 0.01, *** *p* < 0.001. All analyses were carried out using custom-made Python scripts^84^.

### Previous use of the data in other work

Some of the recordings included in the present study have also been used for previous work12, where the specificity of subthreshold responses for the familiar and novel environments was analyzed.

## Acknowledgments

We thank Jérôme Epsztein for his insightful comments on the manuscript and Lucile Le Chevalier-Sontag and Claire Lecestre for their technical assistance. This work was supported by grants from the European Research Council (StG 678790 NEWRON to C.S.-H.), the Pasteur Weizmann Council, the École Doctorale Cerveau-Cognition-Comportement (ED3C, ED n°158, contrat doctoral n°2802/2017 to R.G.-O.) and the Human Frontier Science Program Organization (RGP 0057/2016 to R.M. and M.T.)

## Author contributions

C.S.-H. conceived the project. C.S.-H. and R.G.-O. designed experiments with input from L.P.. R.M., S.C. and M.T. conceived computational modeling. R.G.-O. performed experiments. M.T. performed computational modeling. R.G.-O. analyzed the experimental data with input from C.S.-H. and L.P.. C.S.-H. supervised the project. All authors wrote the manuscript.

## Supplementary information

**Fig. S1.**
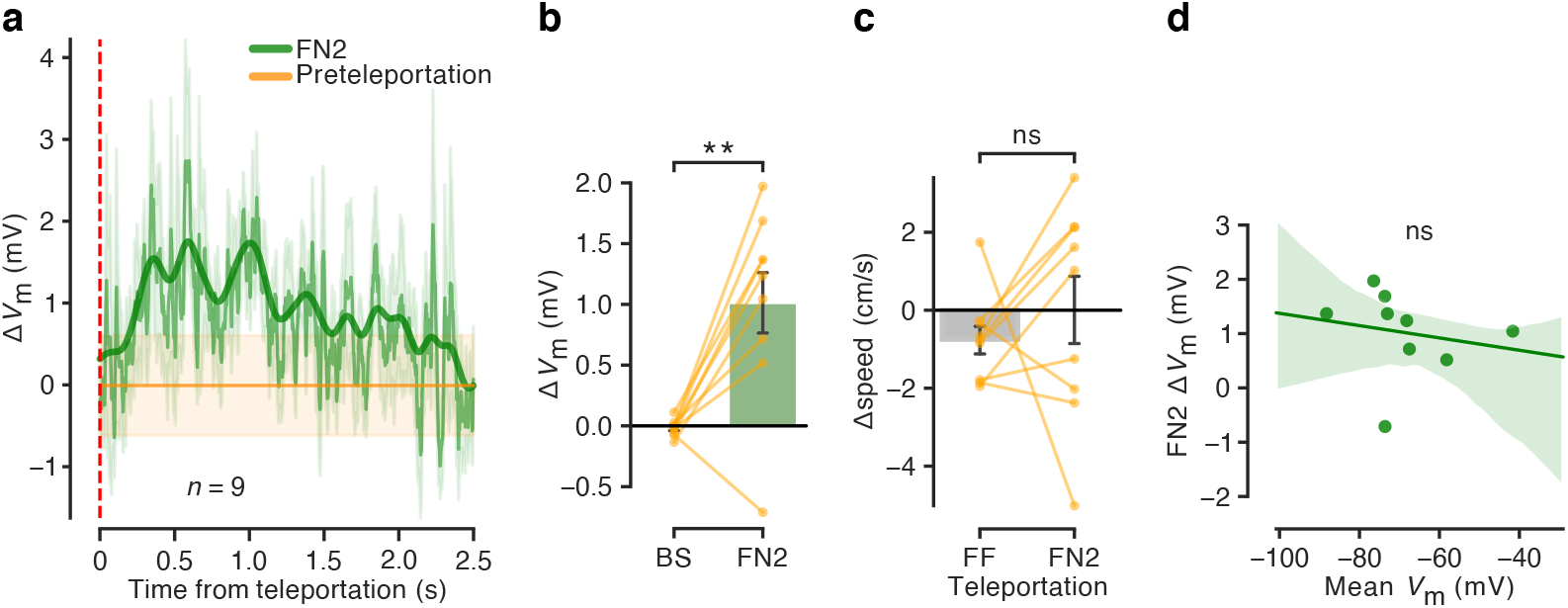
Supplementary analysis to Fig. 2. **a** Teleportation-aligned average across multiple recordings from granule cells showing the temporal dynamics of the subthreshold depolarization in response to novelty. The continuous green trace represents the mean ± s.e.m. of the Δ*V*_m_ recorded 2.5 s after the FN2 teleportation events. The continuous orange trace represents the average mean ± s.e.m. of a 1 s period preceding the teleportation event. Teleportation time is indicated by the vertical red dashed line. A low-pass filtered trace (bold trace) is shown superimposed. **b** Δ*V*_m_ summary for FN2 teleportations tested against a bootstrap obtained from the same dataset (Bootstrap: −0.02 ± 0.02 mV, FN2: 1.02 ± 0.25 mV; *n* = 9 cells, paired t-test, *p* < 0.01). Right bar: same data as in Fig. 2e middle, right bar. **c**Δspeed summary for FF and FN2 teleportations (−0.8 ± 0.4 cm/s and 0.0 ± 0.9 cm/s, respectively; *n* = 9 cells, paired t-test, *p* > 0.05). **d** Correlation between Mean *V*_m_ and FN2 Δ*V*_m_ (*n* = 9 cells, Pearson’s correlation coefficient, *r* = −0.19, *p* > 0.05).

**Fig. S2.**
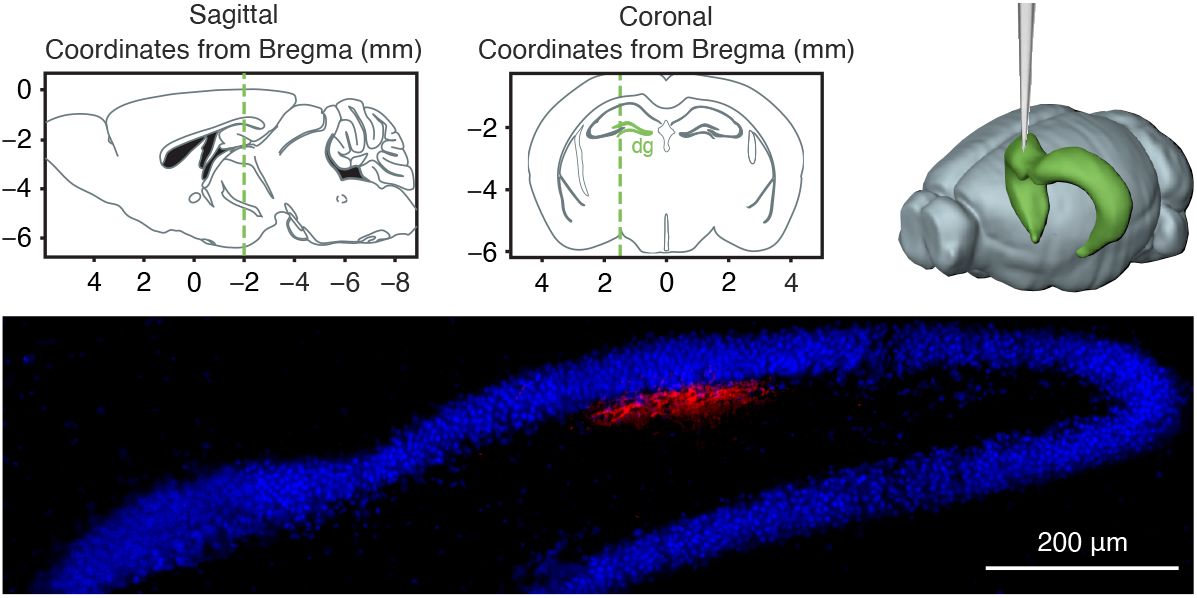
Targeted stereotaxic injections in the dentate gyrus. Coordinates used to target the dentate gyrus: from bregma, anteroposterior −2.0 mm, parasagittal +1.5 mm, depth from cortical surface 1.7 mm. Top left: sagittal view. Top middle: coronal view. Top right: 3D schematic of the target injection site. Bottom: representative example of an injection of the fluorescent marker BODIPY (red) at the target coordinates. DAPI (blue) was used as a nuclear staining to reveal the general anatomy of the preparation. Images generated using the Allen Institute Brain Explorer 2 software (http://mouse.brain-map.org/static/brainexplorer) and Paxinos and Franklin’s The Mouse Brain in Stereotaxic Coordinates.

**Fig. S3.**
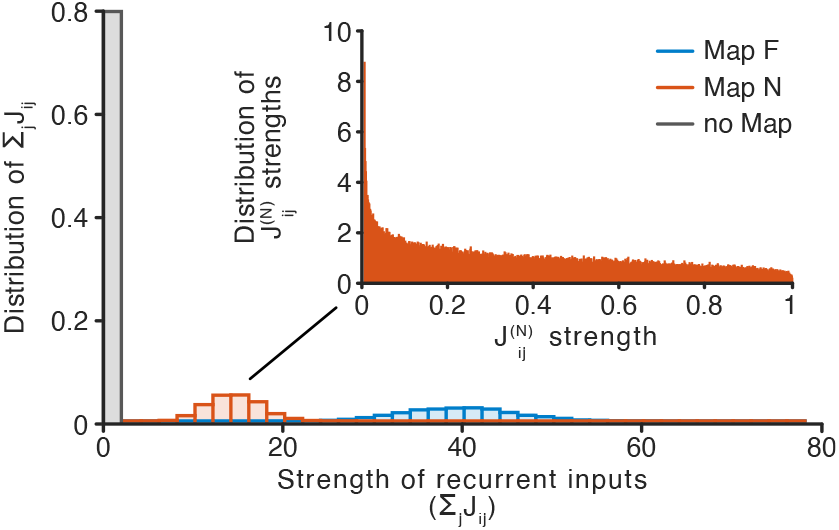
Distribution of synaptic weights in the CA3 network model shows separate bumps for the two maps. Histograms of the summed synaptic inputs for units in map F (blue), units in pre-map N (red) and for the remaining units (grey). Distributions are calculated separately for the two maps with a fraction of units per map of 0.2*n*. Sparser and weaker connections for pre-map N result in a lower average synaptic input to its units compared to the subnetwork storing the F map. Insert plot: histogram (normalized as a probability density function) for modified weights for connections in the N map following a beta distribution with parameters *α* = 0.7, *β* = 1.2. Map N is initially generated as a consolidated map similarly to map F, non-zero weights are then replaced by the beta distributed random variable in the plot. Notice the peak of the distribution at 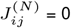.

**Fig. S4.**
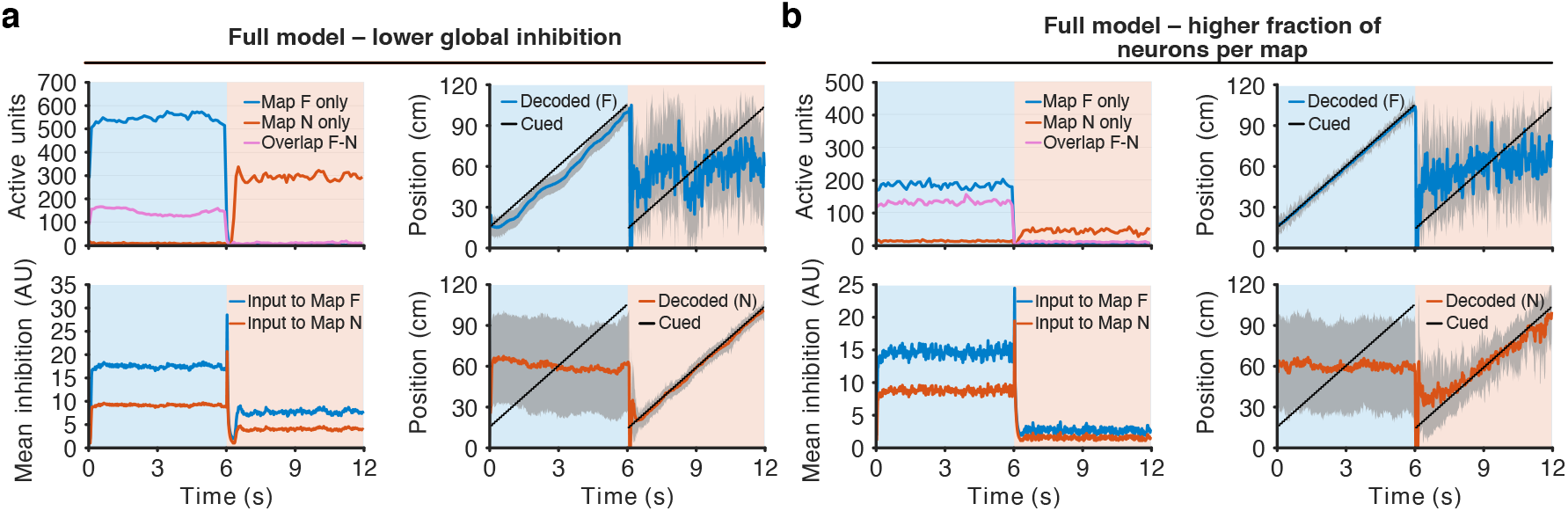
Simulations with alternative sets of network parameters. Effect of global inhibition and fraction of units per map on network dynamics, represented as in Fig. 5b-e. **a** Model with lower coefficients of global inhibition. Activity (top left) is higher in both maps compared to the model in Fig. 5b with no significant differences in network performance. The stronger bump in map F is associated with stronger recurrent inputs, changing the balance with mEC inputs and effectively slowing the bump from following the animal position in the familiar environment (top right). To balance the difference in inhibition in map N, the threshold of initial activation of the map, *S*min, has to be lowered accordingly. This new threshold, affecting also map F, requires a corresponding decrease of the coefficient of global inhibition *g_i,F_* also for this map. Finally, the strength of the transient feed-forward inhibition needs to be adapted to the new levels of inhibition in the network to ensure the disruption of the activity bump in map F. The parameters for this simulation, modified from Fig. 5, were *g_i,N_* = 0.01, *g_i,F_* = 0.025, *S*_min_ = 80 and *A*^inter^ = 30. Similar but opposite variations were also tested (i.e., *g_i,N_* = 0.02, *g_i,F_* = 0.05, *S*_min_ = 20 and *A*^inter^ = 15) with, again, no significant differences in network dynamics. **b** Model with a higher fraction (0.4*n*) of units per map. The increased map size with a fixed number of units per activity pattern generates less correlated memories in the network. Lower correlations between the activity patterns require stronger inhibition and mEC inputs for map F to drive its activity bump, resulting in a lower activity in the F map (top left, blue line) compared to Fig. 5b-e. Activity in the F-N overlap units (top left, purple line) is instead compensated by the increased number of units shared by the two subnetworks. The higher inhibition for units in the F map together with the higher overlap result in a reduced activity (top left, red line) and a noisier spatial selectivity (bottom right) for the N map. The parameters modified from Fig. 5 were *g_i,F_* = 0.045, *S*_min_ = 45, *A*^mEC,F^ = 5 and *A*^inter^ = 30.

**Supplementary Video** Network activity during two simulated lap crossings across the familiar (F) and the novel (N) environment. Blue and red bars represent the number of active cells with a place field center in the corresponding 5 cm bin in, respectively, the F (top) and N (middle) maps. Black dashed lines represent the position cued through the mEC input to the subnetworks involved in the encoding of the two maps. Simulated animal position is shown at the bottom. Note the activity bump moving from the F to the N map after teleportation.

## References

1. Hasselmo, M. E., Schnell, E. & Barkai, E. Dynamics of learning and recall at excitatory recurrent synapses and cholinergic modulation in rat hippocampal region CA3. J Neurosci 15, 5249–5262 (1995).

2. Rolls, E. T. & Treves, A. Neural Networks and Brain Function (Oxford University Press, Oxford, 1998).

3. Frankland, P. W. & Bontempi, B. The organization of recent and remote memories. Nat Rev Neurosci 6, 119–130 (2005).

4. Rolls, E. T. The storage and recall of memories in the hippocampo-cortical system. Cell Tissue Res 373, 577–604 (2018).

5. Wills, T. J., Lever, C., Cacucci, F., Burgess, N. & O’Keefe, J. Attractor dynamics in the hippocampal representation of the local environment. Science 308, 873–876 (2005).

6. Leutgeb, S., Leutgeb, J. K., Moser, M.-B. & Moser, E. I. Place cells, spatial maps and the population code for memory. Curr Opin Neurobiol 15, 738–746 (2005).

7. Neunuebel, J. P. & Knierim, J. J. CA3 retrieves coherent representations from degraded input: direct evidence for CA3 pattern completion and dentate gyrus pattern separation. Neuron 81, 416–427 (2014).

8. Knierim, J. J. & Neunuebel, J. P. Tracking the flow of hippocampal computation: Pattern separation, pattern completion, and attractor dynamics. Neurobiol Learn Mem 129, 38–49 (2016).

9. Guzman, S. J., Schlögl, A., Frotscher, M. & Jonas, P. Synaptic mechanisms of pattern completion in the hippocampal CA3 network. Science 353, 1117–1123 (2016).

10. Hainmueller, T. & Bartos, M. Parallel emergence of stable and dynamic memory engrams in the hippocampus. Nature 558, 292–296 (2018).

11. Hainmueller, T. & Bartos, M. Dentate gyrus circuits for encoding, retrieval and discrimination of episodic memories. Nat Rev Neurosci 21, 153–168 (2020).

12. Allegra, M., Posani, L., Gómez-Ocádiz, R. & Schmidt-Hieber, C. Differential Relation between Neuronal and Behavioral Discrimination during Hippocampal Memory Encoding. Neuron 108, 1103–1112.e6 (2020).

13. Marr, D. A theory of cerebellar cortex. J Physiol 202, 437–470 (1969).

14. Marr, D. Simple memory: a theory for archicortex. Philos Trans R Soc Lond B Biol Sci 262, 23–81 (1971).

15. Albus, J. S. A theory of cerebellar function. Mathematical Biosciences 10, 25–61 (1971).

16. Gilbert, P. E., Kesner, R. P. & Lee, I. Dissociating hippocampal subregions: double dissociation between dentate gyrus and CA1. Hippocampus 11, 626–636 (2001).

17. Hargreaves, E. L., Rao, G., Lee, I. & Knierim, J. J. Major dissociation between medial and lateral entorhinal input to dorsal hippocampus. Science 308, 1792–1794 (2005).

18. Leutgeb, J. K., Leutgeb, S., Moser, M.-B. & Moser, E. I. Pattern separation in the dentate gyrus and CA3 of the hippocampus. Science 315, 961–966 (2007).

19. Diamantaki, M., Frey, M., Berens, P., Preston-Ferrer, P. & Burgalossi, A. Sparse activity of identified dentate granule cells during spatial exploration. Elife 5 (2016).

20. Cayco-Gajic, N. A. & Silver, R. A. Re-evaluating Circuit Mechanisms Underlying Pattern Separation. Neuron 101, 584–602 (2019).

21. Lee, H., GoodSmith, D. & Knierim, J. J. Parallel processing streams in the hippocampus. Curr Opin Neurobiol 64, 127–134 (2020).

22. Kafkas, A. & Montaldi, D. How do memory systems detect and respond to novelty? Neurosci Lett 680, 60–68 (2018).

23. Gava, G. P. et al. Integrating new memories into the hippocampal network activity space. Nat Neurosci 24, 326–330 (2021).

24. Prince, L. Y., Bacon, T. J., Tigaret, C. M. & Mellor, J. R. Neuromodulation of the Feedforward Dentate Gyrus-CA3 Microcircuit. Front Synaptic Neurosci 8, 32 (2016).

25. Palacios-Filardo, J. & Mellor, J. R. Neuromodulation of hippocampal long-term synaptic plasticity. Curr Opin Neurobiol 54, 37–43 (2019).

26. Chen, S. et al. A hypothalamic novelty signal modulates hippocampal memory. Nature 586, 270–274 (2020).

27. Zhang, X., Schlögl, A. & Jonas, P. Selective Routing of Spatial Information Flow from Input to Output in Hippocampal Granule Cells. Neuron 107, 1212–1225.e7 (2020).

28. Bourboulou, R. et al. Dynamic control of hippocampal spatial coding resolution by local visual cues. Elife 8 (2019).

29. Schmidt-Hieber, C. & Nolan, M. F. Synaptic integrative mechanisms for spatial cognition. Nat Neurosci 20, 1483–1492 (2017).

30. Leal, S. L. & Yassa, M. A. Integrating new findings and examining clinical applications of pattern separation. Nat Neurosci 21, 163–173 (2018).

31. Vandael, D., Borges-Merjane, C., Zhang, X. & Jonas, P. Short-Term Plasticity at Hippocampal Mossy Fiber Synapses Is Induced by Natural Activity Patterns and Associated with Vesicle Pool Engram Formation. Neuron 107, 509–521.e7 (2020).

32. Yu, A. J. & Dayan, P. Uncertainty, neuromodulation, and attention. Neuron 46, 681–692 (2005).

33. Teles-Grilo Ruivo, L. M. & Mellor, J. R. Cholinergic modulation of hippocampal network function. Front Synaptic Neurosci 5, 2 (2013).

34. Dannenberg, H., Young, K. & Hasselmo, M. E. Modulation of Hippocampal Circuits by Muscarinic and Nicotinic Receptors. Front Neural Circuits 11, 102 (2017).

35. Samsonovich, A. & McNaughton, B. L. Path integration and cognitive mapping in a continuous attractor neural network model. J Neurosci 17, 5900–5920 (1997).

36. Trouche, S. et al. Recoding a cocaine-place memory engram to a neutral engram in the hippocampus. Nat Neurosci 19, 564–567 (2016).

37. Mori, M., Abegg, M. H., Gähwiler, B. H. & Gerber, U. A frequency-dependent switch from inhibition to excitation in a hippocampal unitary circuit. Nature 431, 453–456 (2004).

38. Zucca, S., Griguoli, M., Malézieux, M., Grosjean, N., Carta, M. & Mulle, C. Control of Spike Transfer at Hippocampal Mossy Fiber Synapses In Vivo by GABAA and GABAB Receptor-Mediated Inhibition. J Neurosci 37, 587–598 (2017).

39. Wilson, M. A. & McNaughton, B. L. Dynamics of the hippocampal ensemble code for space. Science 261, 1055–1058 (1993).

40. Marder, E. Neuromodulation of neuronal circuits: back to the future. Neuron 76, 1–11 (2012).

41. Kandel, E. R., Dudai, Y. & Mayford, M. R. The molecular and systems biology of memory. Cell 157, 163–186 (2014).

42. Hasselmo, M. E. The role of acetylcholine in learning and memory. Curr Opin Neurobiol 16, 710–715 (2006).

43. Hasselmo, M. E. Neuromodulation: acetylcholine and memory consolidation. Trends Cogn Sci 3, 351–359 (1999).

44. Aloisi, A. M., Casamenti, F., Scali, C., Pepeu, G. & Carli, G. Effects of novelty, pain and stress on hippocampal extracellular acetylcholine levels in male rats. Brain Res 748, 219–226 (1997).

45. Ceccarelli, I., Casamenti, F., Massafra, C., Pepeu, G., Scali, C. & Aloisi, A. M. Effects of novelty and pain on behavior and hippocampal extracellular ACh levels in male and female rats. Brain Res 815, 169–176 (1999).

46. Giovannini, M. G., Rakovska, A., Benton, R. S., Pazzagli, M., Bianchi, L. & Pepeu, G. Effects of novelty and habituation on acetylcholine, GABA, and glutamate release from the frontal cortex and hippocampus of freely moving rats. Neuroscience 106, 43–53 (2001).

47. Bianchi, L., Ballini, C., Colivicchi, M. A., Della Corte, L., Giovannini, M. G. & Pepeu, G. Investigation on acetylcholine, aspartate, glutamate and GABA extracellular levels from ventral hippocampus during repeated exploratory activity in the rat. Neurochem Res 28, 565–573 (2003).

48. Blokland, A., Honig, W. & Raaijmakers, W. G. Effects of intra-hippocampal scopolamine injections in a repeated spatial acquisition task in the rat. Psychopharmacology (Berl) 109, 373–376 (1992).

49. Ohno, M., Yamamoto, T. & Watanabe, S. Blockade of hippocampal nicotinic receptors impairs working memory but not reference memory in rats. Pharmacol Biochem Behav 45, 89–93 (1993).

50. Ohno, M., Yamamoto, T. & Watanabe, S. Blockade of hippocampal M1 muscarinic receptors impairs working memory performance of rats. Brain Res 650, 260–266 (1994).

51. Rogers, J. L. & Kesner, R. P. Cholinergic modulation of the hippocampus during encoding and retrieval. Neurobiol Learn Mem 80, 332–342 (2003).

52. Pabst, M. et al. Astrocyte Intermediaries of Septal Cholinergic Modulation in the Hippocampus. Neuron 90, 853–865 (2016).

53. Overstreet Wadiche, L., Bromberg, D. A., Bensen, A. L. & Westbrook, G. L. GABAergic signaling to newborn neurons in dentate gyrus. J Neurophysiol 94, 4528–4532 (2005).

54. Ge, S., Goh, E. L. K., Sailor, K. A., Kitabatake, Y., Ming, G.-l. & Song, H. GABA regulates synaptic integration of newly generated neurons in the adult brain. Nature 439, 589–593 (2006).

55. Richards, C. D. Anaesthetic modulation of synaptic transmission in the mammalian CNS. Br J Anaesth 89, 79–90 (2002).

56. Hao, X. et al. The Effects of General Anesthetics on Synaptic Transmission. Curr Neuropharmacol 18, 936–965 (2020).

57. Nitz, D. & McNaughton, B. Differential modulation of CA1 and dentate gyrus interneurons during exploration of novel environments. J Neurophysiol 91, 863–872 (2004).

58. Wess, J. Novel insights into muscarinic acetylcholine receptor function using gene targeting technology. Trends Pharmacol Sci 24, 414–420 (2003).

59. Levey, A. I., Edmunds, S. M., Koliatsos, V., Wiley, R. G. & Heilman, C. J. Expression of m1-m4 muscarinic acetylcholine receptor proteins in rat hippocampus and regulation by cholinergic innervation. J Neurosci 15, 4077–4092 (1995).

60. Yamasaki, M., Matsui, M. & Watanabe, M. Preferential localization of muscarinic M1 receptor on dendritic shaft and spine of cortical pyramidal cells and its anatomical evidence for volume transmission. J Neurosci 30, 4408–4418 (2010).

61. Losonczy, A., Makara, J. K. & Magee, J. C. Compartmentalized dendritic plasticity and input feature storage in neurons. Nature 452, 436–441 (2008).

62. Petrovic, M. M., Nowacki, J., Olivo, V., Tsaneva-Atanasova, K., Randall, A. D. & Mellor, J. R. Inhibition of post-synaptic Kv7/KCNQ/M channels facilitates long-term potentiation in the hippocampus. PLoS One 7, e30402 (2012).

63. Giessel, A. J. & Sabatini, B. L. M1 muscarinic receptors boost synaptic potentials and calcium influx in dendritic spines by inhibiting postsynaptic SK channels. Neuron 68, 936–947 (2010).

64. Buchanan, K. A., Petrovic, M. M., Chamberlain, S. E. L., Marrion, N. V. & Mellor, J. R. Facilitation of long-term potentiation by muscarinic M(1) receptors is mediated by inhibition of SK channels. Neuron 68, 948–963 (2010).

65. Hofmann, M. E. & Frazier, C. J. Muscarinic receptor activation modulates the excitability of hilar mossy cells through the induction of an afterdepolarization. Brain Res 1318, 42–51 (2010).

66. Anderson, R. W. & Strowbridge, B. W. Regulation of persistent activity in hippocampal mossy cells by inhibitory synaptic potentials. Learn Mem 21, 263–271 (2014).

67. Fredes, F., Silva, M. A., Koppensteiner, P., Kobayashi, K., Joesch, M. & Shigemoto, R. Ventro-dorsal Hippocampal Pathway Gates Novelty-Induced Contextual Memory Formation. Curr Biol 31, 25–38.e5 (2021).

68. Diamantaki, M., Frey, M., Preston-Ferrer, P. & Burgalossi, A. Priming Spatial Activity by Single-Cell Stimulation in the Dentate Gyrus of Freely Moving Rats. Curr Biol 26, 536–541 (2016).

69. Huang, L. et al. Relationship between simultaneously recorded spiking activity and fluorescence signal in GCaMP6 transgenic mice. Elife 10(2021).

70. Steinmetz, N. A., Koch, C., Harris, K. D. & Carandini, M. Challenges and opportunities for large-scale electrophysiology with Neuropixels probes. Curr Opin Neurobiol 50, 92–100 (2018).

71. Vyleta, N. P., Borges-Merjane, C. & Jonas, P. Plasticity-dependent, full detonation at hippocampal mossy fiber-CA3 pyramidal neuron synapses. Elife 5(2016).

72. Dragoi, G. & Tonegawa, S. Preplay of future place cell sequences by hippocampal cellular assemblies. Nature 469, 397–401 (2011).

73. Posani, L., Cocco, S. & Monasson, R. Integration and multiplexing of positional and contextual information by the hippocampal network. PLoS Comput Biol 14, e1006320 (2018).

74. Sheffield, M. E. J., Adoff, M. D. & Dombeck, D. A. Increased Prevalence of Calcium Transients across the Dendritic Arbor during Place Field Formation. Neuron 96, 490–504.e5 (2017).

75. Dupret, D., O’Neill, J. & Csicsvari, J. Dynamic reconfiguration of hippocampal interneuron circuits during spatial learning. Neuron 78, 166–180 (2013).

76. Barak, O., Rigotti, M. & Fusi, S. The sparseness of mixed selectivity neurons controls the generalization-discrimination trade-off. J Neurosci 33, 3844–3856 (2013).

77. Schmidt-Hieber, C. & Häusser, M. Cellular mechanisms of spatial navigation in the medial entorhinal cortex. Nat Neurosci 16, 325–331 (2013).

78. Schmidt-Hieber, C. et al. Active dendritic integration as a mechanism for robust and precise grid cell firing. Nat Neurosci 20, 1114–1121 (2017).

79. Robinson, N. T. M. et al. Targeted Activation of Hippocampal Place Cells Drives Memory-Guided Spatial Behavior. Cell 183, 1586–1599.e10 (2020).

80. Guo, Z. V. et al. Procedures for behavioral experiments in head-fixed mice. PLoS One 9, e88678 (2014).

81. Paxinos, G. & Franklin, K. B. J. Paxinos and Franklin’s the Mouse Brain in Stereotaxic Coordinates, Compact 5th Edition (Elsevier Academic Press, San Diego, 2019).

82. Margrie, T. W., Brecht, M. & Sakmann, B. In vivo, low-resistance, whole-cell recordings from neurons in the anaesthetized and awake mammalian brain. Pflugers Arch 444, 491–498 (2002).

83. Monasson, R. & Rosay, S. Crosstalk and transitions between multiple spatial maps in an attractor neural network model of the hippocampus: collective motion of the activity. Phys Rev E Stat Nonlin Soft Matter Phys 89, 032803 (2014).

84. Harris, C. R. et al. Array programming with NumPy. Nature 585, 357–362 (2020).

